# Lis1 binding regulates force-induced detachment of cytoplasmic dynein from microtubules

**DOI:** 10.1101/2022.06.02.494578

**Authors:** Emre Kusakci, Zaw Min Htet, Yuanchang Zhao, John P. Gillies, Samara L. Reck-Peterson, Ahmet Yildiz

## Abstract

Cytoplasmic dynein-1 (dynein) is an AAA+ motor that transports intracellular cargos towards the microtubule minus end. Lissencephaly-1 (Lis1) binds to the AAA+ ring and stalk of dynein’s motor domain and promotes the assembly of active dynein complexes. Recent studies showed that Lis1 slows motility when it remains bound to dynein, but the underlying mechanism remained unclear. Using single-molecule and optical trapping assays, we investigated how Lis1 binding affects the motility and force generation of yeast dynein in vitro. We showed that Lis1 does not slow dynein motility by serving as a roadblock or tethering dynein to microtubules. Lis1 binding also does not affect the forces that stall dynein movement, but it induces prolonged stalls and reduces the asymmetry in the force-induced detachment of dynein from microtubules. The mutagenesis of the Lis1 binding sites on dynein’s stalk partially recovers this asymmetry but does not restore dynein velocity. These results suggest that Lis1’s interaction with the AAA+ ring is sufficient to result in slower movement and that Lis1’s interaction with dynein’s stalk slows force-induced detachment of dynein from microtubules.

---

Cytoplasmic dynein-1 (dynein hereafter) is the primary motor responsible for motility and force generation towards the minus ends of microtubules^1^. Dynein transports membranous organelles, vesicles, mRNA, and unfolded proteins towards the nucleus, drives retrograde transport in neurons, and plays crucial roles in cell division^2^. Mutations that impair dynein function are linked to severe neurodegenerative and developmental disorders^3^.

The core structural component of the dynein transport machinery is the dynein heavy chain (DHC), which is composed of the tail domain and motor domain (Fig. 1a). The tail domain facilitates dimerization and recruits other associated chains. The motor domain forms a catalytic AAA+ ring that connects to the tail via the linker domain and to the microtubule via a coiled-coil stalk^4, 5^ (Fig. 1a). ATP hydrolysis in the AAA+ ring is coupled to the swinging motion of the linker at the surface of the ring^1^, which powers minus-end-directed motility^6^. Nucleotide hydrolysis also controls the binding and release of the motor from the microtubule by altering the registry of the stalk coiled coils^7^. Isolated dynein adopts an autoinhibited “phi” conformation and exhibits diffusive or nonprocessive motility on microtubules^8-10^. Processive motility of mammalian dynein is activated when it forms a complex with dynactin and the coiled-coil domain of a cargo adaptor protein^11, 12^.

**Figure 1.**
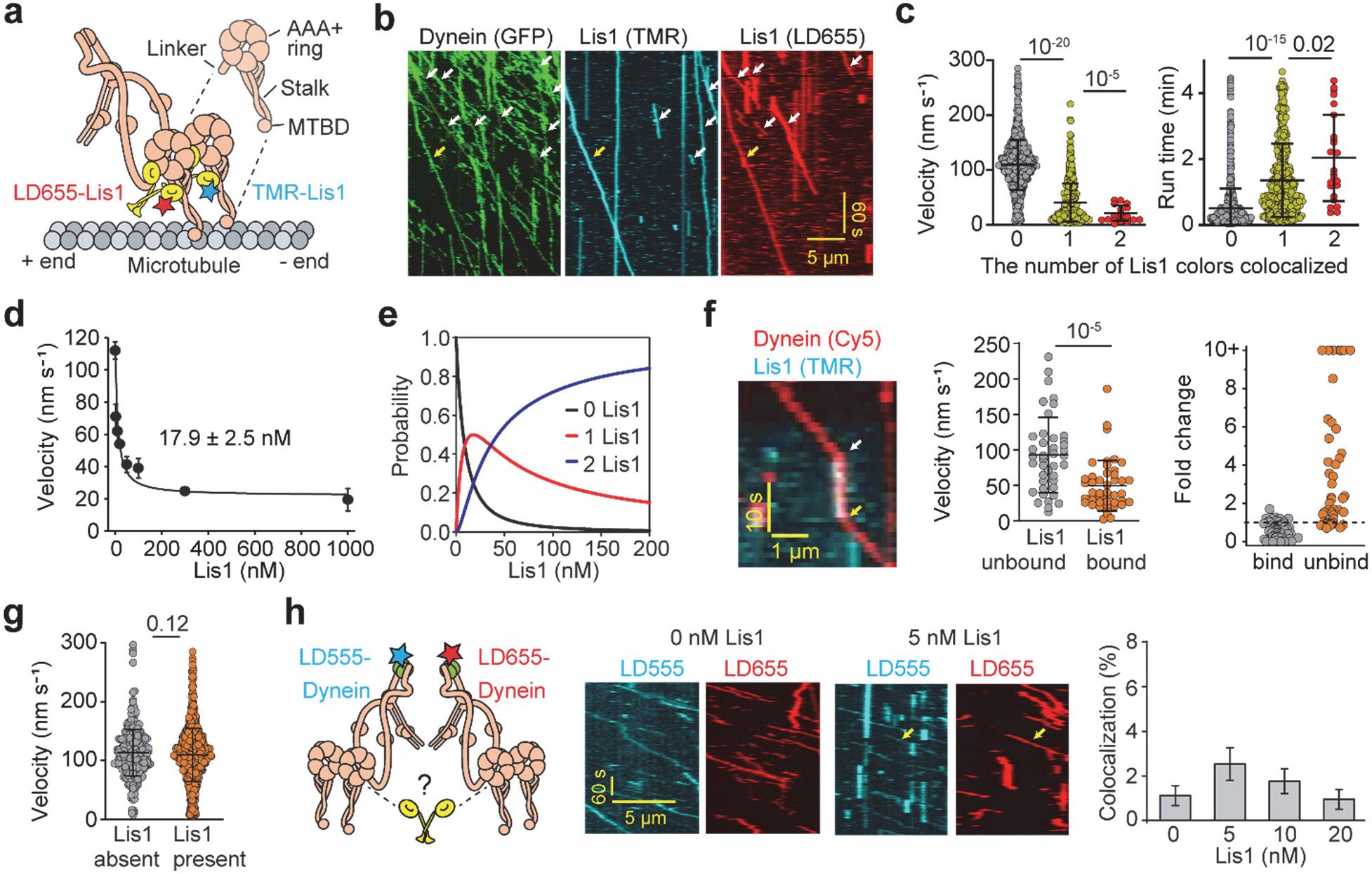
Lis1 binding reduces the velocity while increasing the microtubule residence time of dynein. **a**) Schematic of the single-molecule motility of yeast cytoplasmic dynein on microtubules in the presence of fluorescently labeled Lis1. Insert shows the structural organization of the dynein motor domain. **b)** A representative kymograph of GFP-dynein in the presence of 2 nM LD655- and 2 nM TMR-labeled Lis1. Arrows highlight processive motors colocalized with one (white) and two (yellow) colors of Lis1. **c)** The velocity and run time of dynein motors colocalized with 0, 1, and 2 colors of Lis1 (mean ± s.d., N = 1706, 671, and 24 from left to right). **d)** Dynein velocity under different Lis1 concentrations in 50 mM KAc (mean ± s.e.m.; N = 861, 534, 233, 352, 278, 312, and 131 for 0,10, 20, 50, 100, 300, and 1,000 nM Lis1, respectively; three biological replicates). The fit of the velocity data (solid curve, see Methods) reveals K_D_ (±s.e.). **e)** The estimated probability of 0, 1, and 2 Lis1 bound to a dynein dimer under different Lis1 concentrations (see Methods). **f)** (Left) A representative trace of transient binding (white arrow) and unbinding (yellow arrow) of Lis1 from dynein during processive motility. (Middle) The velocity of Lis1-unbound and Lis1-bound sections of individual dynein trajectories (mean ± s.d., N = 41 and 45 from left to right). (Right) The fold change in motor velocity upon binding or unbinding of Lis1 (mean ± s.d., N = 64 and 47 from left to right). The velocity of the motor before the binding or unbinding event was normalized to 1.0. **g)** The velocities of individual dynein motors in the absence of Lis1 compared to the motors that do not colocalize with Lis1 when fluorescently labeled Lis1 is present in the chamber (N = 361 and 1534, from left to right). **h)** (Left) Schematic shows whether a single Lis1 dimer can crossbridge two dynein dimers labeled with different dyes. (Middle) Example kymographs of 5 nM LD555-dynein (cyan) and 5 nM LD655-dynein (red) in 0 and 5 nM Lis1. Yellow arrows show colocalization between LD555 and LD655 dyes. (Right) The percentage of dynein trajectories that exhibit colocalization between LD555 and LD655 dyes (mean ± s.d., N = 408, 378, 426, and 208 from left to right, three independent experiments). In **c**, **f**, and **g**, the centerline and whiskers represent the mean and s.d., respectively. P-values were calculated by a two-tailed t-test with Welch correction for velocity and by Kolmogorov-Smirnov test for run time.

Dynein-mediated transport also requires Lis1, which is the only known regulatory protein that directly binds to the dynein motor domain^13^. Heterozygous mutations in the *LIS1* gene lead to neuronal migration deficiency during embryonic brain development and cause the severe neurodevelopmental disease, lissencephaly^14^. Studies in live cells showed that Lis1 is a required cofactor for the recruitment of dynein to kinetochores, the nuclear envelope, and the cell cortex, and the initiation of dynein-mediated transport of a wide variety of cargos, including Rab6-vesicles, lysosomes, and ribonucleoprotein complexes (reviewed in^15^). Lis1 consists of an amino-terminal dimerization domain and a carboxy-terminal WD40 β-propeller domain. These β-propeller domains of a Lis1 dimer bind to dynein’s motor domain near the AAA3 site of the AAA+ ring and the base of the stalk (Fig. 1a)^13, 16, 17^. Recent in vitro and in vivo studies have proposed that Lis1 binding to the AAA+ ring facilitates the assembly of active dynein-dynactin-adaptor complexes by preventing dynein from adopting the phi conformation^18-21^. Structure-guided mutations showed that the Lis1-stalk interaction is important for the localization of dynein to the cortex of yeast cells^16, 17^, but it remains to be determined how this interaction affects the mechanism of dynein motility.

The comigration of Lis1 has been reported to substantially reduce the velocity of microtubule gliding or processive motility of yeast, pig, and human dynein-dynactin in vitro^13, 18, 19, 22-24^. Several models have been proposed to explain how Lis1 binding pauses or slows down dynein motility. Lis1 binding increases dynein’s microtubule-binding affinity^13, 22, 23, 25^, which may result in slower detachment of the motor domain before it can step along the microtubule. Cryo-electron microscopy of Lis1-bound dynein suggested that Lis1 clamps multiple AAA subunits together, trapping dynein in a conformation with high microtubule binding affinity^16^, possibly preventing rearrangements of the AAA subunits needed to switch to the low-affinity conformation. In addition, Lis1 binding to dynein has been proposed to inhibit or slow dynein by blocking the powerstroke of its linker domain^25^. A recent in vitro study proposed that Pac1 (the Lis1 homolog in *S. cerevisiae*, “Lis1” hereafter) reduces dynein velocity in low ionic strength buffers by simultaneously interacting with the microtubule and exerting a drag by tethering dynein to the microtubule^20^. There are also reports that the Lis1 association increases^26^ or does not affect^27^ the velocity of human dynein-dynactin. Therefore, how Lis1 affects dynein velocity remains controversial.

In this study, we used single-molecule imaging and optical trapping to investigate how Lis1 binding affects the motility and force generation of *S. cerevisiae* cytoplasmic dynein. Similar to mammalian dynein, Lis1 binding to yeast dynein prevents it from adopting the phi confirmation, and dynein mutants that cannot form the phi conformation partially rescue dynein function in yeast lacking Lis1^20^. We previously showed that yeast dynein has similar stepping and force generation properties to mammalian dynein-dynactin-adaptor complexes^28-31^. Although yeast dynein also requires dynactin and a cortical attachment protein for its cellular function^32, 33^, it can walk processively in the absence of these cofactors in vitro^34^. Therefore, yeast dynein serves as a simpler model to investigate how Lis1 binding affects the intrinsic motility and force generation properties of dynein. Consistent with previous observations^13, 18, 19, 35^, here we find that Lis1 binding slows dynein motility and induces longer run times on microtubules. Lis1 weakly interacts with microtubules, but this interaction does not slow dynein motility because Lis1 binding slows dynein even under conditions in which Lis1 does not interact with microtubules. Optical trapping assays revealed that Lis1 does not reduce the dynein stall force, but results in slower microtubule detachment of dynein under hindering forces. Lis1 also decreases the asymmetry in the force-induced velocity of dynein. Mutations that disrupt Lis1’s interactions with dynein’s stalk partially restore the asymmetric detachment of dynein from microtubules in the presence of Lis1. These observations provide new insight into the mechanism of dynein regulation by Lis1.

## Results

### Lis1 binding stoichiometrically slows dynein motility

We expressed the full-length DHC (*DYN1*) with an amino-terminal GFP tag and a carboxy-terminal HaloTag from its endogenous locus in an *S. cerevisiae* strain lacking the genes encoding Lis1 (*PAC1*), the Lis binding protein, NudEL (*NDL1*), and the p150 subunit of dynactin (*NIP100*, Extended Data Table 1). The DHC co-purified with the endogenous dynein light intermediate chain (Dyn3), light chain (Dyn2), and intermediate chain (Pac11)^34^. We monitored the motility of this endogenously expressed complex (hereafter dynein) on surface-immobilized microtubules in the presence and absence of Lis1 using multi-color total-internal reflection fluorescence (TIRF) microscopy (Fig. 1a). To determine how Lis1 binding affects dynein motility, we separately labeled two different batches of Lis1 with TMR and LD655 dyes and monitored the colocalization of 2 nM TMR-Lis1 and 2 nM LD655-Lis1 with GFP-dynein. In this assay, colocalization of dynein with a single color of Lis1 might be due to one or two Lis1 dimers bound per dynein, whereas colocalization of both dyes with the GFP signal ensures that two Lis1 dimers are bound to a dynein dimer, one on each motor domain. We observed either LD655-Lis1 or TMR-Lis1 translocating together with dynein on microtubules at similar colocalization percentages (14% each; Fig. 1b,c, Supplementary Movie 1). A small fraction (1%) of GFP-dynein motors colocalized with LD655- and TMR-Lis1 simultaneously, suggesting that a single dynein motor can recruit two Lis1 dimers (Fig. 1b), but this occurred rarely at low (2 nM) Lis1 concentrations^27^. Similar to mammalian dynein-dynactin-adaptor complexes^18, 19^, yeast dynein motors that colocalized with a single color of Lis1 moved slower (41 nm s^-1^) and had a longer run time (81.4 s) on average than motors that did not colocalize with Lis1 (110 nm s^-1^ velocity and 30.5 s run time). Colocalization of both Lis1 colors further reduced the velocity (21 nm s^-1^) and increased the run time (122.5 s) of dynein (Fig. 1c).

To estimate the affinity of Lis1 binding to processive dynein motors, we monitored the motility of dynein motors in the presence of 0-250 nM unlabeled Lis1 (Extended Data Fig. 1a-c). The average velocity of dynein under different Lis1 concentrations was fitted to the fraction of motors bound to 0, 1, and 2 Lis1s multiplied with their corresponding average velocities (Fig. 1d, see Methods). The fit estimated that the dissociation constant of Lis1 (K_D_) is 17.9 nM (Fig. 1d) and that most of the motors colocalize with two Lis1s at high Lis1 concentrations (Fig. 1e). We also observed that transient binding of Lis1 paused or slowed 83% of the motors, whereas unbinding of Lis1 increased the velocity of 77% of the motors, in agreement with previous observations on mammalian dynein^27^ (Fig. 1f, Extended Data Fig. 1d, Supplementary Movie 2). The velocity of dynein motors not colocalizing with Lis1 was similar to dynein velocity when there was no Lis1 in the chamber (Fig. 1g). Therefore, Lis1 slows dynein motility only when it is directly bound to the motor, and the presence of excess Lis1 in the chamber does not affect dynein velocity.

Previously, we proposed that human Lis1 facilitates the recruitment of two dyneins to the dynactin-adaptor complex by opening dynein’s autoinhibited phi-conformation^18, 19^. However, it is also possible that β-propeller domains of a Lis1 dimer cross-bridge the motor domains of two separate dynein dimers (Fig. 1i) and recruit them together to the dynactin-adaptor complex. To test this possibility, we differentially labeled two batches of dynein with LD555 and LD655 dyes and monitored their comigration on microtubules in the presence or absence of Lis1. Because the yeast strain we purify endogenous dynein from lacks the *NIP100* gene, a functional dynactin complex should not be present in our assays, and colocalization of two dyneins could only be mediated by Lis1. In the absence of Lis1, we observed less than 1% colocalization between dynein labeled with equal concentrations of LD555 and LD655. The presence of equimolar Lis1 to dynein led to only a marginal increase (2%) in colocalization between the differentially labeled dyneins (Fig. 1i), suggesting that Lis1 does not effectively cross-bridge two dyneins.

### Lis1’s interaction with the microtubule does not affect dynein velocity

Unlike mammalian Lis1^18, 19^, yeast Lis1 interacts with microtubules^20^. The degree to which Lis1 reduces dynein velocity was reported to correlate with the extent of Lis1–microtubule binding under different salt concentrations, indicating that yeast Lis1 may slow dynein by simultaneously binding dynein and the microtubule^20^. However, it is also possible that increasing the ionic strength of the buffer weakens the microtubule affinity of dynein. This could compete against the Lis1-mediated increase in the microtubule affinity of the motor, thereby reducing the effect of Lis1 in altering dynein velocity. We envisioned that the microtubule tethering model requires that 1) either one Lis1 monomer interacts with dynein and the other monomer interacts with the microtubule, or 2) a single Lis1 monomer interacts simultaneously with both dynein and the microtubule (Fig. 2a). To test these models, we first characterized microtubule binding dynamics of wild-type Lis1 dimer (Lis1^WT^) and monomeric Lis1 lacking its amino-terminal dimerization domain (Lis1^monomer^)^13^. As previously reported^20^, Lis1^WT^ decorated microtubules in 50 mM KAc (Fig. 2b,c). The fit to a binding isotherm revealed a K_D_ of 88 ± 8 nM (Extended Data Fig. 2a), higher than the K_D_ value we calculated from the Lis-mediated reduction of dynein velocity (Fig. 1d). Single Lis1^WT^ stayed bound to the microtubule for 19.3 s on average and exhibited diffusive dynamics on the microtubule surface (Fig. 2d,e). However, there was little to no microtubule decoration of Lis1^WT^ when the ionic strength of the buffer was increased to a physiologically relevant level (150 mM KAc; Fig. 2f,g)^20^ or the carboxy-terminal tails of tubulin were cleaved by subtilisin ^20^ (Fig. 2h, Extended Data Fig. 2b). We concluded that Lis1 interacts with negatively charged tails of tubulin primarily through electrostatic interactions. However, this interaction is not strong enough to sustain microtubule binding in physiologically relevant salt concentrations, consistent with the lack of microtubule colocalization of Lis1 in yeast^36^. Compared to Lis1^WT^, Lis1^monomer^ exhibits little to no microtubule binding even under low salt conditions. Lis1^monomer^ had a lower affinity for microtubules (Fig. 2b,c), only transiently interacted with microtubules, and typically dissociated within 1-2 s (Fig. 2d,e).

**Figure 2.**
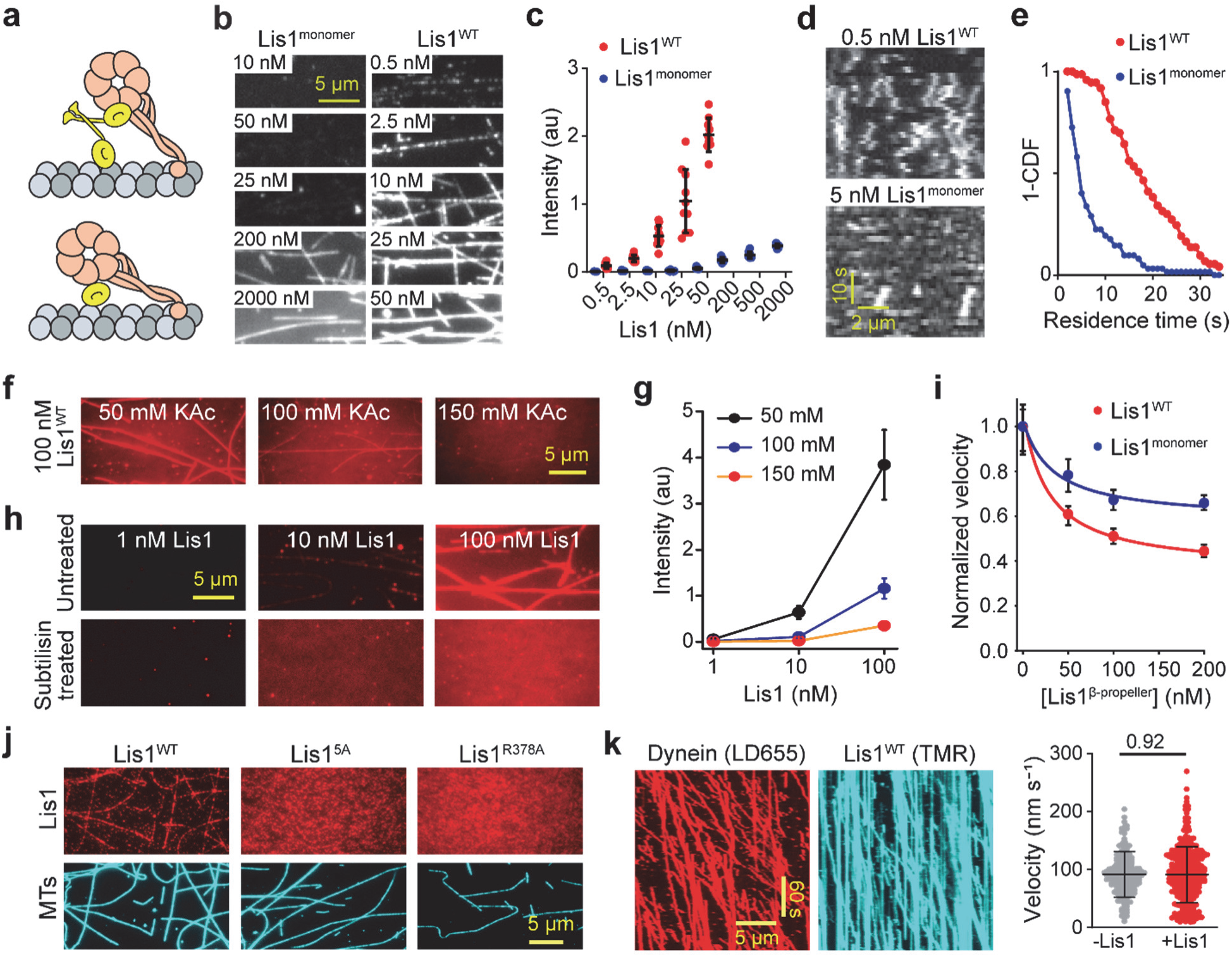
Lis1’s interaction with microtubules does not slow down dynein motility. **a)** Models of dynein tethered to a microtubule either via one Lis1 β-propeller docked to dynein and the other on the microtubule (top) or one Lis1 β-propeller simultaneously interacts with dynein and the microtubule (bottom). **b)** Representative images of different concentrations of LD655-labeled Lis1^WT^ and Lis1^monomer^ binding to surface-immobilized microtubules. **c)** Fluorescence intensities of Lis1^WT^ and Lis1^monomer^ on microtubules (mean ± s.d., N = 10 microtubules per condition; three technical replicates). **d)** Representative kymographs show landing, diffusion, and dissociation of Lis1^WT^ and Lis1^monomer^ from microtubules. **e)** The inverse cumulative distribution function (1-CDF) of the microtubule residence time of Lis1^WT^ and Lis1^monomer^ (N = 73 for Lis1^WT^ and 62 for Lis1^monomer^). Molecules that reside on microtubules less than 2 frames (2 s) were excluded from the analysis. **f)** Lis1^WT^ binding to microtubules is reduced by increasing the salt concentration. **g)** The average intensity of fluorescently labeled Lis1^WT^ on microtubules under different salt concentrations (mean ± s.d., N = 10 microtubules per condition; three technical replicates). **h)** Lis1^WT^ does not decorate subtilisin-treated microtubules in 50 mM KAc. **i)** Single-molecule velocity of dynein with increasing concentrations of Lis1^WT^ or Lis1^monomer^. The median and interquartile ranges are shown. Data were normalized to a velocity in the absence of Lis1 (N = 400 runs per condition). Solid curves represent a fit to reveal the K_D_ (28 nM for Lis1^WT^ and 35 nM for Lis1^monomer^) and the minimum value for the normalized velocity (0.36 for Lis1^WT^ and 0.58 for Lis1^monomer^). **j**) Two-color imaging shows that, unlike Lis1^WT^, Lis1^5A^ and Lis1^R378A^ do not bind microtubules. Lis1 concentration was set to 100 nM and the assays were performed in 50 mM KAc. **k)** (Left) A representative kymograph of dynein motility on Lis1-decorated microtubules. Occasionally, dynein picks up Lis1 on microtubules and comigrates with it during processive motility. Lis1 trajectories without comigrating dynein are due to incomplete labeling of the motor. (Right) Velocities of dynein motors on undecorated (-Lis1) and Lis1-decorated (+Lis1) microtubules. (N= 208 and 329, from left to right). The P-value was calculated by a two-tailed t-test with Welch correction. In **c** and **k**, the center line and whiskers represent the mean and s.d., respectively.

We next measured the velocity of dynein in the presence of either Lis1^WT^ or Lis1^monomer^. The first model predicts that Lis1^monomer^ would not affect dynein’s velocity. However, consistent with our previous observations^13^, Lis1^monomer^ slowed dynein motility in a dose-dependent manner (Fig. 2i), albeit not as strongly as Lis1^WT^. Thus, if Lis1 crosslinks dynein to microtubules, it must do so in the context of a monomer, with one face of Lis1 interacting with dynein and a different one interacting with microtubules. The second model predicts that mutants that disrupt Lis1 binding to dynein would not disrupt Lis1 microtubule binding as the interfaces involved must be different. We tested this possibility by using dynein binding mutants of Lis1, in which a single (Lis1^R378A^) or five (Lis1^5A^) positively charged residues in Lis1’s β-propeller domain were mutated to alanine (Extended Data Fig. 2c)^25^. Unlike Lis1^WT^, Lis1^R378A^ and Lis1^5A^ were unable to decorate microtubules in 50 mM KAc (Fig. 2j, Extended Data Fig. 2d-e). Lis1^5A^ also did not bind to microtubules in microtubule pelleting assays (Extended Data Fig. 2a), indicating that Lis1 interacts with microtubules through basic amino acids at the dynein-interacting surface of its β-propeller domain. Because a Lis1^monomer^ would not be capable of crosslinking dynein to microtubules, our results are inconsistent with the microtubule tethering model.

While our results show that Lis1 binding to dynein is required to slow dynein motility (Fig. 1), Marzo et al. reported that the presence of Lis1 is sufficient to reduce the velocity of dynein motors that do not comigrate with Lis1^20^. This observation raises the possibility that Lis1 serves as a static obstacle against dynein motility on the microtubule surface, akin to microtubule-associated proteins (MAPs)^37, 38^. Because all dyneins would encounter microtubule bound Lis1, this model predicts a similar reduction in velocity for dyneins that comigrate with Lis1 and those that are not bound to Lis1^20^. To test this model, we pre-decorated microtubules with 300 nM Lis1 with no added salt and washed away free Lis1 from the chamber (Extended Data Fig. 2f). We found that 92% of Lis1 intensity disappeared after washing the chamber and introducing dynein, consistent with transient and diffusive interactions of Lis1 with the microtubule. The fraction of Lis1 remaining on the microtubules failed to impede dynein motility (Fig. 2k, Supplementary Movies 3 and 4), indicating that Lis1 does not serve as an effective roadblock on the microtubule against dynein motility.

We also purified dynein from the *S. cerevisiae* strain used by Marzo et al., which overexpresses DHC and each of its associated chains under the galactose promoter (hereafter dynein_Gal_)^20^. Similar to dynein expressed using the endogenous promoter (Fig. 1), dynein_Gal_ had an increased run time when colocalized with Lis1 (Extended Data Figs. 3-4). However, dynein_Gal_ motors moved significantly slower whether or not they colocalized with Lis1 in the chamber (Extended Data Fig. 4d,f), as reported^20^. The reduction in dynein_Gal_ velocity was not due to the decoration of the microtubule surface by Lis1, because Lis1 addition slowed dynein_Gal_ even when it did not efficiently decorate microtubules under physiological salt (150 mM; Extended Data Fig. 3). We next introduced or removed Lis1 from the chamber while recording dynein_Gal_ motility. Introducing Lis1 into the chamber did not alter the velocity of dynein_Gal_ motors that were already moving along the microtubule at the time Lis1 was added (Extended Data Fig. 5a-c). Likewise, the removal of free Lis1 from the chamber failed to recover dynein_Gal_ velocity (Extended Data Fig. 5c), indicating that dynein_Gal_ velocity is reduced when it is preincubated with Lis1 before it is bound to the microtubule, not by the comigration of Lis1 with the motor. While the underlying mechanism remains unclear, the differences between the Lis1-mediated regulation of dynein and dynein_Gal_ may be related to low stability or aggregation of dynein_Gal_ after purification^20^ (see Methods).

### Lis1 does not affect the dynein stall force

We next investigated whether Lis1 binding slows dynein motility by interfering with the swinging motion of the linker at the surface of the ring^25^. Because the linker drives the force-generating powerstroke of dynein, this model predicts that Lis1 binding reduces the ability of dynein to walk against hindering forces. To test this possibility, we measured the stall force of endogenously expressed dynein in the presence and absence of excess (300 nM) Lis1 using an optical trap (Fig. 3a). In the absence of Lis1, dynein stalled at 3.8 ± 0.1 pN (mean ± s.e.m., N = 116 stalls) and remained attached to the microtubule for 14.6 ± 0. 8 s (mean ± s.e.m.) before the trapped bead snapped back to the trap center^31^ (Fig. 3b,c). The stall force was unaltered by Lis1 addition (3.8 ± 0.1 pN, N = 126 stalls; Fig. 3b,c), suggesting that Lis1 does not disrupt the force-generating powerstroke of the linker. However, the Lis1 addition resulted in a 70% increase in the duration of the stalls (24.7 ± 0.9 s, p = 0.018, Kolmogorov-Smirnov test), indicating that Lis1 reduces the microtubule detachment rate of dynein under hindering forces (Fig. 3d)^18, 35^.

**Figure 3.**
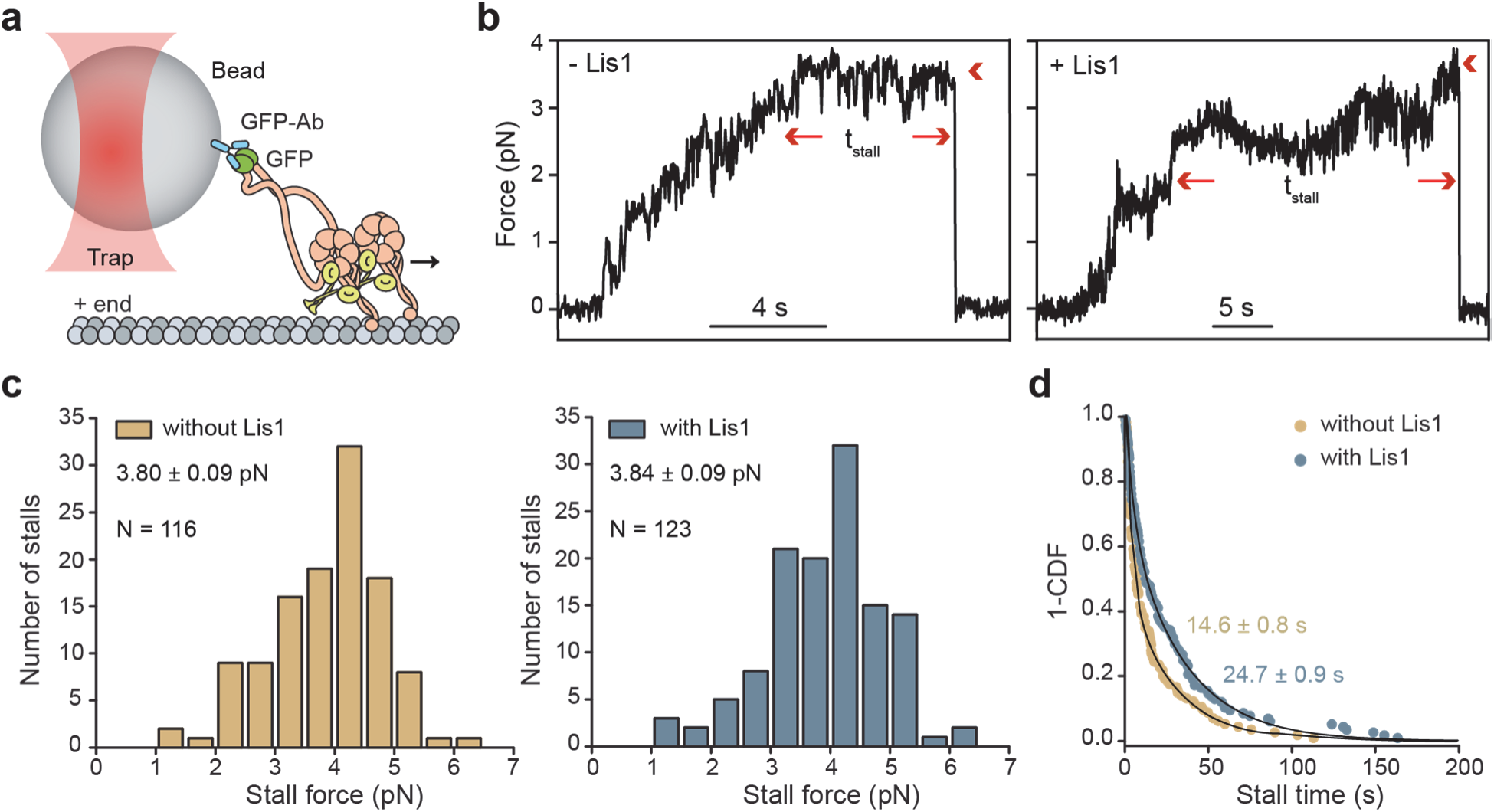
Lis1 does not affect the stall force of dynein. **a)** The schematic shows stall force measurements of GFP-dynein in the presence of unlabeled Lis1 using a fixed-beam optical trap. **b)** Sample trajectories of beads driven by single dynein motors in 1 mM ATP with and without 300 nM Lis1. The arrows show the beginning and the end of a stalling event. The arrowheads indicate the detachment of the motor from the microtubule and the snapping of the bead to the trap center. Stall time (t_stall_) was defined as the time the bead spends at the 30% margin of the stall force before it detaches from the microtubule. **c)** The stall force histogram of dynein in the presence and absence of 300 nM Lis1 (mean ±s.e.m.; p = 0.77, two-tailed t-test). **d)** Stall times of dynein with and without 300 nM Lis1. Fit to a double exponential decay (solid curves) reveals the weighted average of t_stall_ (±s.e.).

Unlike full-length dynein, Lis1 binding reduced the stall force of tail-truncated and artificially dimerized dynein constructs^31, 39^, suggesting that artificial dimerization of the linkers at the exit of the ring introduces a steric obstacle against their swinging motion when Lis1 is present on the outer surface of the ring (Extended Data Fig. 6). However, Lis1 binding does not affect the force generation of full-length dynein, presumably because the longer length of the full-length dynein tails has greater flexibility compared to the truncated dynein constructs used by Toropova et al.^25^.

### Lis1 reduces the asymmetry in force-induced detachment of dynein

Because Lis1 increases the microtubule affinity of dynein (Fig. 1)^13, 22, 23, 25^, we also tested whether Lis1 binding slows motility by altering the detachment kinetics of dynein from the microtubule. Dynein controls its microtubule affinity by altering the registry of the stalk coiled coils through nucleotide-dependent conformational changes in the AAA+ ring^5^. The stalk sliding mechanism is also sensitive to external forces^40^. When pulled in the assisting direction (towards the minus-end of microtubules), the stalk switches to a weak-binding registry, and dynein releases quickly from the microtubule and moves substantially faster. However, the stalk remains in a strongly bound registry and the motor resists backward movement when pulled in the hindering direction^31, 40-43^.

If Lis1 binding interferes with the stalk sliding mechanism, we anticipated the addition of Lis1 to alter the force-velocity (F-V) behavior of dynein. To test this prediction, we first measured the velocity of single dynein motors in 1 mM ATP and in the absence of Lis1 when they were subjected to constant forces in assisting and hindering directions using a force-feedback controlled trap. Consistent with previous reports ^31, 39, 43^, hindering force slowed minus-end directed motility of dynein and the motors start walking towards the plus-end under superstall forces. In comparison, the minus-end directed velocity of dynein motility increased more rapidly under the same magnitude forces in the assisting direction (Fig. 4a). The addition of 300 nM Lis1 reduced dynein speed when the motor was being pulled in either direction (Fig. 4b), but the speed was reduced more substantially under assisting forces than under hindering forces (Fig. 4c,d). To quantify the changes in the asymmetry in F-V of dynein, we compared the speed of the motor when it walks towards the plus-end versus the minus end of the microtubule under force. Because dynein changes its direction relative to the stall force, not by the actual direction of the applied force, in the presence of ATP^31, 39^, we defined the asymmetry as the ratio of forward and backward speeds of dynein when the motor is subjected to the same magnitude of forces relative to the stall force (F_rel_). Consistent with our hypothesis, Lis1 addition substantially reduced the asymmetry between plus- and minus-end directed speeds of dynein from 2.45 to 1.14, on average (Fig. 4c,d).

**Figure 4.**
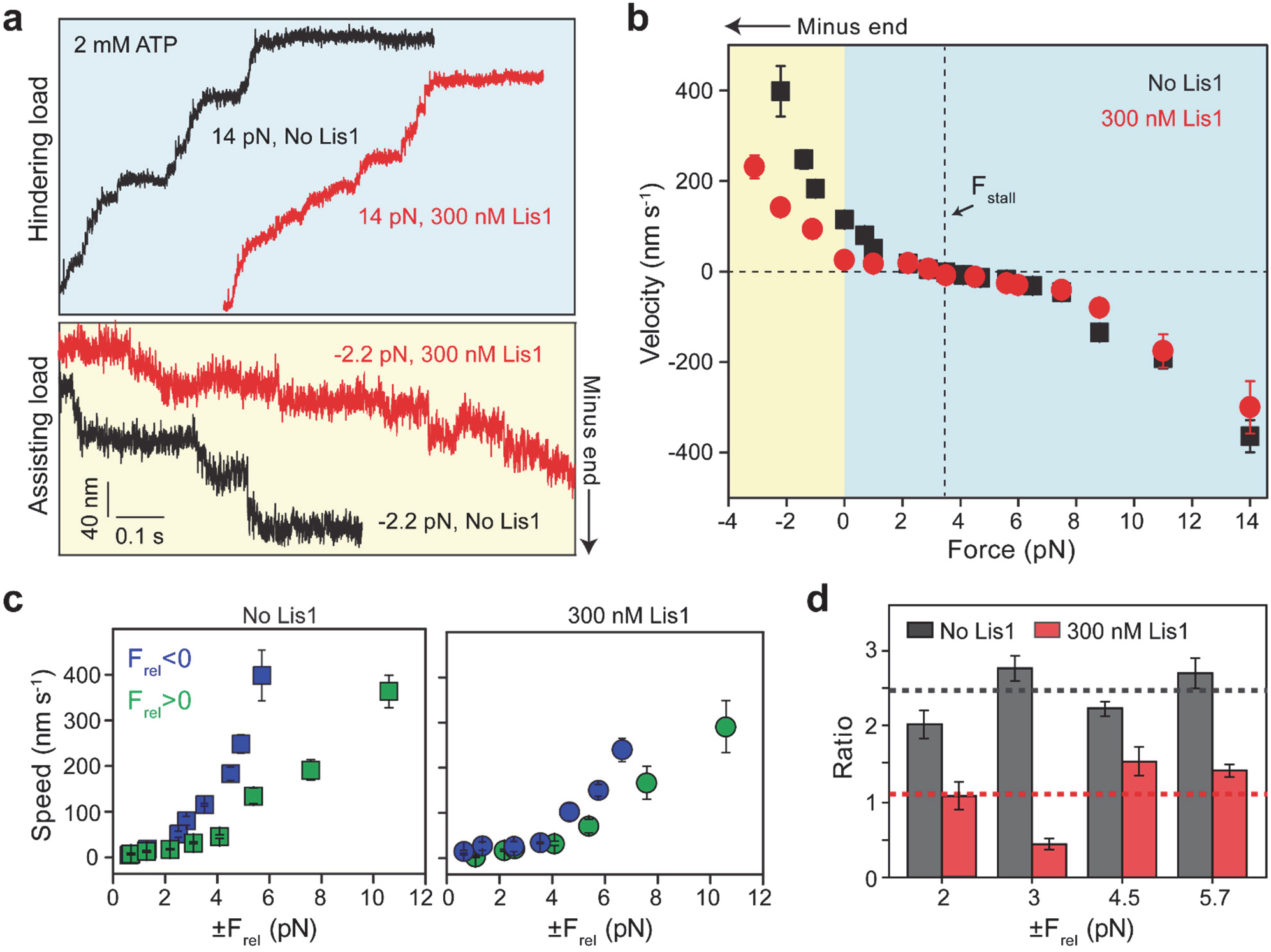
Lis1 reduces the asymmetry in the F-V behavior of dynein in the presence of ATP. **a)** Representative traces of dynein-driven beads under hindering (positive) and assisting (negative) forces with and without 300 nM Lis1. Assays were performed in 2 mM ATP and 50 mM KAc. **b)** F-V measurements of dynein in the presence and absence of 300 nM Lis1 (mean ±s.e.m., from left to right, N = 41, 38, 37, 861, 27, 52, 106, 28, 51, 60, 89, 115, 124, 62, 23, 38, and 40 runs without Lis1 and 34, 52, 25, 312, 33, 42, 49, 37, 62, 47, 33, 42, 36, 29, and 18 runs with Lis1). **c)** The speed of dynein-driven beads (mean ± s.e.m.) under the same magnitude of positive and negative forces relative to the stall force (F_rel_). **d)** The ratios of the dynein speeds under the same magnitude of positive and negative F_rel_. The dashed lines represent the average of the asymmetry ratios measured under different forces. The error bars represent s.e.m.

We reasoned that Lis1 binding may reduce the asymmetry in F-V by interfering with either the linker swing or the stalk sliding mechanisms of dynein. The swinging motion of the linker is strictly coupled to the nucleotide-dependent rigid body motions of the AAA+ ring^44^, whereas the registry of the stalk coiled coils can be altered by external force even in the absence of ATP^40^. To distinguish between these possibilities, we tested whether Lis1 still affects the asymmetry in the F-V of dynein in the absence of ATP. Similar to the ATP condition, we observed Lis1 addition to decrease dynein speed when the motor was pulled in both directions and the decrease in speed was more substantial when the motor was pulled in the assisting direction in the absence of ATP (Fig. 5a,b). Because the motor moves in a direction it is pulled in the absence of ATP, we calculated the asymmetry relative to the no-force condition in this case. Lis1 addition reduced the asymmetry from 1.83 to 1.10, on average (Fig. 5c). These results indicate that Lis1 binding slows the detachment of dynein from microtubules by interfering with the stalk sliding mechanism, not by interfering with the swinging motion of the linker.

**Figure 5.**
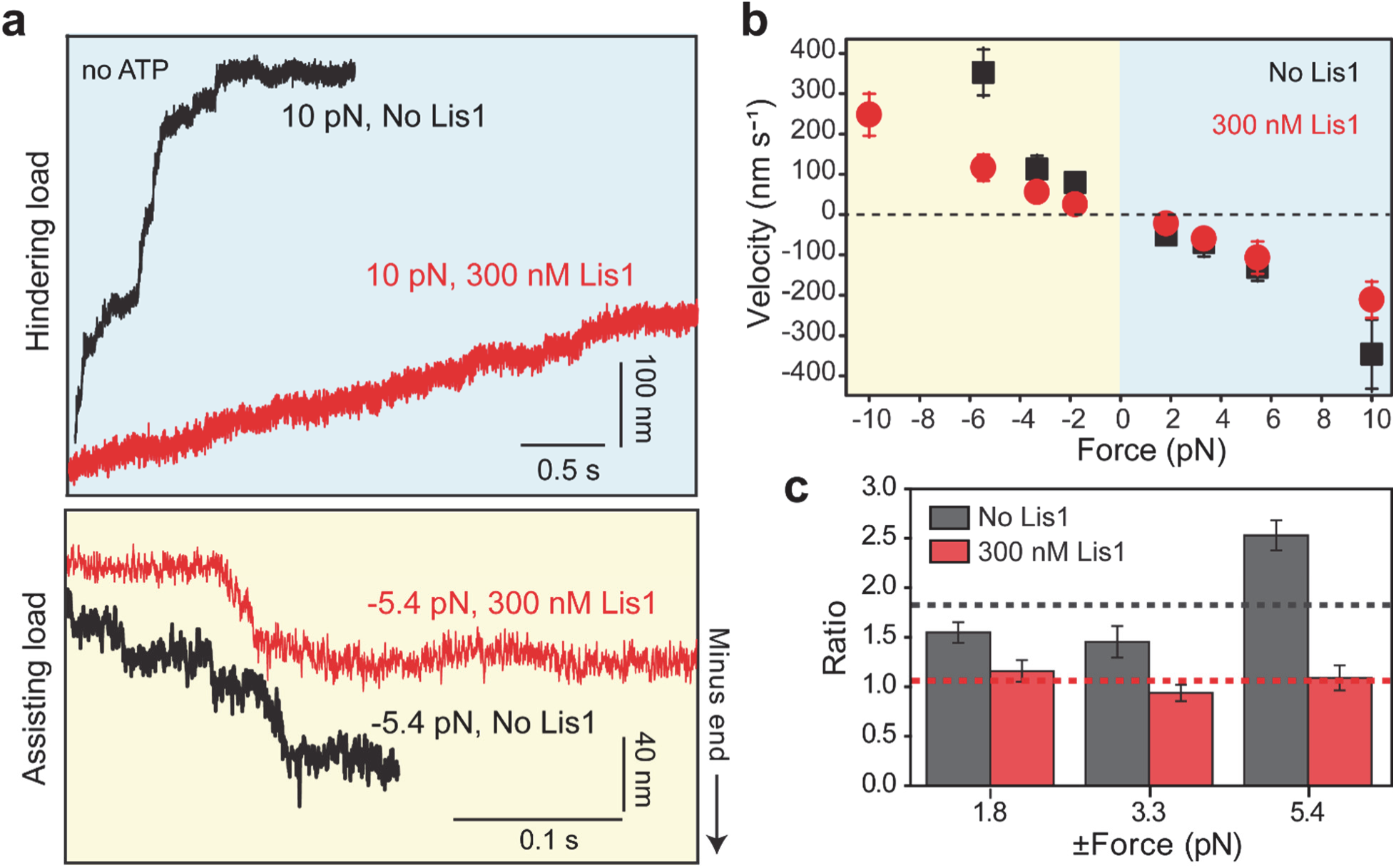
Lis1 reduces the asymmetry in F-V of dynein in the absence of ATP. **a)** Representative traces of dynein-driven beads under assisting and hindering forces with and without 300 nM Lis1. Assays were performed in 50 mM KAc and without ATP. **b)** F-V measurements of dynein in the absence and presence of 300 nM Lis1 (mean ± s.e.m.; from left to right, N = 42, 29, 65, 51, 38, 62, and 15 runs without Lis1 and 36, 35, 34, 28, 21, 25, 20, and 19 runs with Lis1). **c)** The ratios of the velocities under the same magnitude of forces in assisting and hindering directions relative to 0 pN force. The dashed lines represent the average of the asymmetry ratios measured under different forces. The error bars represent s.e.m.

We previously showed that Lis1 stably interacts with a second binding site at the base of dynein’s stalk^17, 45^. To test whether this Lis1-stalk interaction might be responsible for Lis1’s ability to reduce the asymmetry in force-induced detachment of dynein from microtubules, we performed F-V measurements using a dynein mutant, in which three Lis1-interacting residues on its stalk were replaced with alanine (dynein^EQN^, purified from yeast endogenously expressing dynein subunits)^17^. We first confirmed that dynein^EQN^ motility is slowed down by Lis1, albeit with a ∼3-fold higher K_D_ (54 ±10 nM, ±s.e.) compared to that for wild-type (WT) dynein^17^. (Extended Data Fig. 7a,b). Similar to WT dynein, the binding of two Lis1 dimers further slowed dynein^EQN^ motility (Extended Data Fig. 7c,d). In the absence of Lis1, dynein^EQN^ also had similar stall force, but lower stall times relative to WT dynein (p = 0.03, Kolmogorov-Smirnov test, Extended Data Fig. 7e-g). To determine how Lis1 affects stall force and the F-V behavior of dynein^EQN^, we used higher concentrations of Lis1 (900 nM) to compensate for the lower affinity of dynein^EQN^ for Lis1^17^. The addition of Lis1 resulted in only minor changes in the stall force and stall time of dynein^EQN^ (Extended Data Fig. 7e-g). However, unlike WT dynein, the addition of Lis1 did not reduce the asymmetry of dynein^EQN^’s F-V behavior (2.1 without Lis1 and 2.0 with Lis1; Fig. 6 and Extended Data Fig. 8). Collectively, these results show that the Lis1-stalk interaction increases Lis1’s affinity to bind dynein and reduces the asymmetry in force-induced detachment of dynein from the microtubule, but the Lis1-ring interaction is sufficient to increase the microtubule affinity and reduce the velocity of dynein.

**Figure 6.**
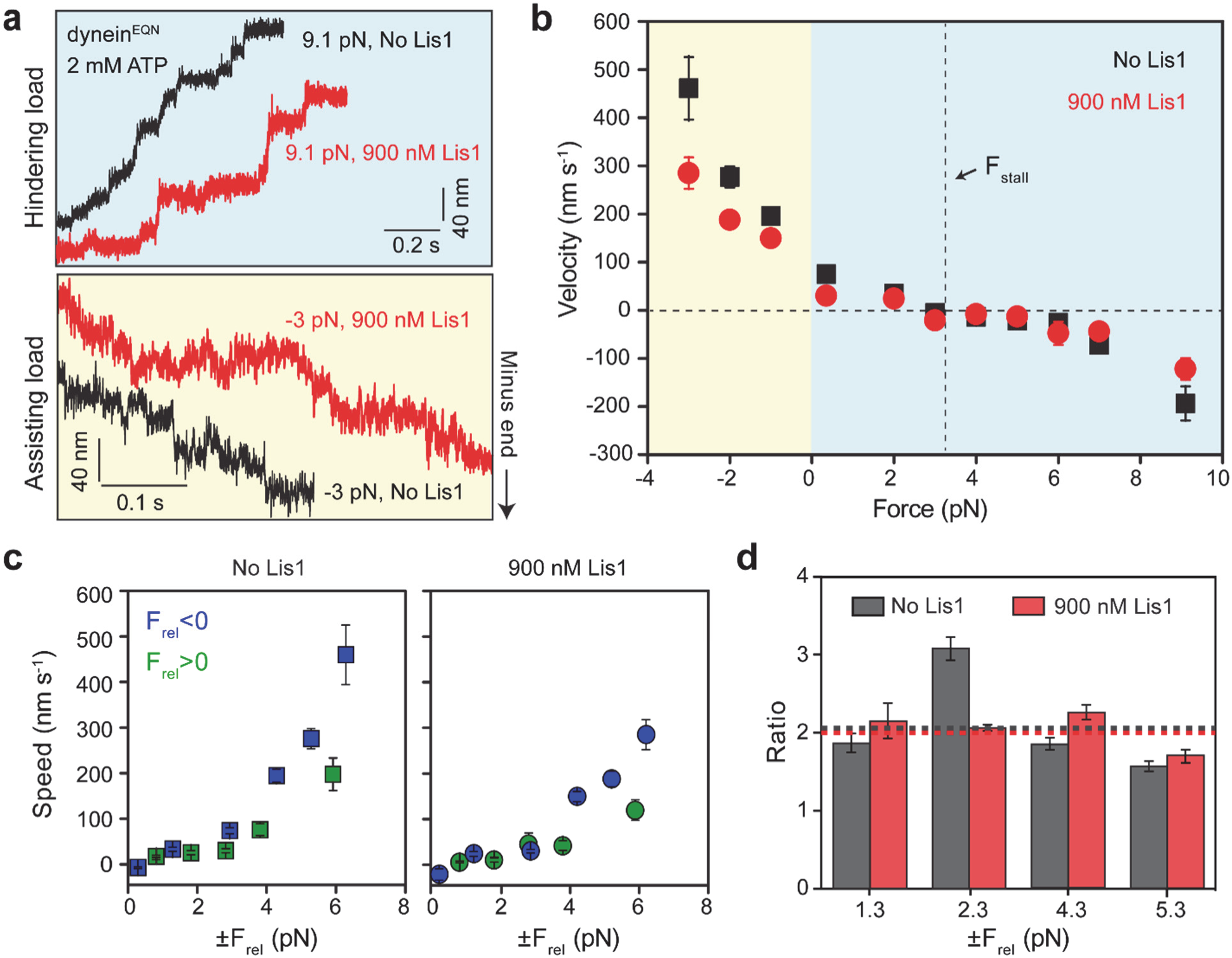
Mutagenesis of the Lis1 binding site on dynein’s stalk recovers the asymmetry in F-V of dynein. **a)** Representative traces of dynein^EQN^-driven beads under assisting and hindering forces with and without 900 nM Lis1. Assays were performed in 2 mM ATP and 50 mM KAc. **b)** F-V measurements of dynein^EQN^ with and without 900 nM Lis1 (mean ± s.e.m.; from left to right, N = 31, 66, 62, 48, 47, 26, 42, 39, 23, 45, and 33 runs without Lis1 and 54, 36, 61, 31, 37, 17, 54, 14, 19, 38, and 26 runs with Lis1). **c)** The speed of dynein^EQN^-driven beads (mean ± s.e.m.) under the same magnitude of positive and negative forces relative to the stall force of dynein^EQN^ (3.3 pN). **d)** The ratios of the velocities under the same magnitude of forces relative to the stall force. The dashed lines represent the average of the asymmetry ratios measured under different forces. The error bars represent s.e.m.

## Discussion

In this study, we used single-molecule imaging to test the models that describe how Lis1 affects the motility of yeast cytoplasmic dynein. Because yeast dynein walks processively in the absence of dynactin and an activating adaptor in vitro, we were able to investigate how Lis1 binding affects the inherent motility and force generation properties of single dynein motors. We observed that when dynein colocalizes with one or two Lis1 dimers, it moves more slowly and runs on microtubules for a longer time than motors that do not colocalize with Lis1. We conclude that Lis1 binding to dynein is necessary to induce increased microtubule affinity and decreased velocity. These results are consistent with our previous observations that Lis1 binding slows the motility of yeast dynein^13, 17^ as well as mammalian dynein-dynactin^18, 19^, and inconsistent with reports that Lis1 colocalization increases^26^ or does not change^27^ the speed of mammalian dynein-dynactin. This discrepancy may be relevant to the two competing effects of Lis1 on the velocity of mammalian dynein-dynactin: Lis1 mediates the assembly of faster complexes that recruit two dyneins to dynactin, but it slows the motility of active complexes if it remains bound to dynein^18, 19^. Therefore, Lis1’s overall effect on mammalian dynein-dynactin might depend on the stoichiometry of Lis1 and dynein in activated dynein-dynactin complexes.

We observed that Lis1 weakly interacts with the C-terminal tails of tubulin at low salt^20^ through the same surface of its β-propeller domain that interacts with dynein. Although its microtubule binding affinity is reduced by nearly two orders of magnitude, Lis1 still effectively slows dynein motility at physiological salt concentrations^20^. In addition, dynein motility can be slowed by a single Lis1 β-propeller domain^13^, which cannot simultaneously interact with dynein and the microtubule. Collectively, our results are inconsistent with a model that Lis1 slows dynein motility by tethering the motor to the microtubule^20^ and show that Lis1 still slows dynein motility under conditions in which it does not interact with the microtubule. Lis1 is also unlikely to serve as an effective roadblock against dynein motility because it exhibits diffusive motion on microtubules and its microtubule decoration does not substantially affect dynein motility along these tracks.

We also performed optical trapping assays to determine how Lis1 binding affects force generation and the force response of dynein. Similar to mammalian dynein^18^, Lis1 binding does not affect the stall force of yeast dynein, indicating that the linker can perform its force-generating powerstroke when Lis1 is bound to the AAA+ ring. This result is consistent with the ability of dynein motors to walk processively, albeit at lower speeds, when colocalized with Lis1^18, 19^, whereas disrupting the linker swing mechanism fully abrogates the motility^46^. Lis1-bound dynein also persists against microtubule detachment for longer durations under hindering forces and moves slower under both assisting and hindering forces, consistent with the high microtubule affinity of Lis1-bound dynein^13, 22, 23, 25^. These results are consistent with previous studies that reported an increase in stall duration of isolated mammalian dynein or the mammalian dynein-dynactin-adaptor complex in the presence of Lis1^18, 35^, underscoring the conservation of Lis1 function across evolution.

Based on our results and previous reports, we propose that Lis1 slows dynein motility in two distinct steps. First, Lis1 binding to the AAA+ ring traps the ring in a conformation that favors high microtubule affinity^17^. Because Lis1 still slows the motility of dynein^EQN^ in the absence of force, Lis1 binding to the AAA+ ring is sufficient for slower movement^16, 17^. Second, Lis1 binding to the stalk traps the coiled coils in a strongly bound registry and restricts their registry shift. In the absence of Lis1, the stalk can switch from the strongly bound to the weakly bound registry when dynein is subjected to assisting forces but remains in a strongly bound registry under hindering forces^40^. Lis1 binding slows dynein speed in both assisting and hindering forces, but it has a more profound effect in limiting the acceleration of dynein under assisting forces. We suggest that this is because Lis1 binding to dynein’s stalk prevents the shift of the coiled coils to a registry with lower microtubule affinity by an external force. Consistent with this model, Lis1 is less effective in reducing dynein speed under assisting forces when the stalk residues that interact with Lis1 were mutated to alanine.

Our results are consistent with Lis1’s role in relieving the autoinhibition of dynein in the phi conformation and facilitating the assembly of the dynein-dynactin complex^18-21^. In vivo studies suggest that Lis1 has additional regulatory roles in addition to rescuing dynein from its autoinhibited conformation, because mutations that disrupt the phi conformation of dynein does not fully rescue loss of endogenous Lis1^20, 21^. Lis1 binding not only opens the phi conformation, but also induces tight microtubule binding of open dynein^35^. This may contribute to the assembly of dynein with dynactin and an activating adaptor by recruiting the motor to the site of assembly and preventing its premature detachment from the microtubule.

Lis1 has been reported to be absent from moving dynein cargos in vivo^47-49^ and it appears to dissociate from most dynein complexes during or after the activation of processive motility in vitro^18-20, 50-52^. Dissociation of Lis1 may be a key step in dynein activation because enhancing the affinity of dynein for Lis1 leads to defects in nuclear migration in the budding yeast, *S. cerevisiae*^36^. However, Lis1 may remain bound to dynein when it transports high load cargos or pulls on astral microtubules during cell division^53, 54^. Lis1 binding may prevent microtubule detachment of dynein when the motor is subjected to not only resistive forces but also to assisting forces during back-and-forth oscillations of spindle microtubules, thereby enabling multiple dyneins to effectively increase tension for proper positioning of the spindle. Future studies are required to determine whether the association and dissociation of Lis1 from dynein are regulated in different cellular contexts.

## Author Contributions

E.K., S.R.P., and A.Y. conceived the study and designed the experiments. Z.M.H. generated the yeast constructs for dynein and Lis1. E.K. purified the proteins. E.K. and Y.Z performed single-molecule experiments and analyzed the data. J.P.G. performed experiments on monomeric Lis1. E.K., S.R.P., and A.Y. wrote the manuscript. All authors read and revised the manuscript.

## Acknowledgments

We thank S. Can, M. ElShenawy, L. S. Ferro, J. T. Canty, and other members of the Yildiz and Reck-Peterson laboratories for helpful discussions, and S. M. Markus for sharing the yeast strain that expresses dynein_Gal_. This work was supported by grants from the National Institute of General Medical Sciences (GM094522, AY), the National Science Foundation (MCB-1617028 and MCB-1055017, AY), and the Fellowship of the Ministry of Education of the Turkish Republic (E.K.). S.R.P. is a Howard Hughes Medical Institute Investigator and is also supported by the National Institutes of Health (1R35GM141825).

## Methods

### Protein purification and labeling

The endogenous genomic copies of *S. cerevisiae* dynein heavy chain (*DYN1*) and Lis1 (*PAC1*) were modified or deleted using homologous recombination^34^. The list of strains used in this study is shown in Extended Data Table 1. The strains that express GFP- and HALO-tagged full-length dynein (RPY1732 and 1736) were generated from the parent yeast strains reported by Reck-Peterson et al.^34^ and Huang et al.^13^. Tagged yeast dynein showed no defects in nuclear segregation (not shown), indicating that dynein’s promoter is not disrupted in this strain background.

*S. cerevisiae* strains that express dynein_Gal_ and Lis1 constructs were grown in 2 L YPA-galactose media (1% yeast extract, 1% peptone, % 0.004 adenine sulfate, 2% galactose), and strains that express endogenously expressed dynein were grown in 2L YPD media (1% yeast extract, 1% peptone, % 0.004 adenine sulfate, 2% dextrose) until OD_600_ reaches 2.0. The cells were pelleted at 6,000 *g* for 15 min, resuspended in phosphate buffer saline (PBS), and frozen in liquid nitrogen. For purification of dynein and dynein_Gal_, cell pellets were ground and dissolved in the dynein lysis buffer (DLB; 30 mM HEPES pH 7.4, 2 mM Mg(Ac)_2_, 1 mM EGTA, 10% glycerol) supplemented with 50 mM KAc, 1 mM DTT, 0.1 mM ATP, 1 mM phenylmethylsulfonyl fluoride (PMSF), 0.1% Triton at 37℃. The lysate was centrifuged at 360,000 *g* for 45 min and the supernatant was incubated with 300 µL IgG beads for 1 h at 4 ℃. The mixture was then applied to Qiagen columns, washed with 30 mL DLB supplemented with 250 mM KAc, 1 mM DTT, 0.1 mM ATP, 1 mM phenylmethylsulfonyl fluoride (PMSF), 0.1% Triton and then with 20 mL TEV buffer (10 mM Tris-HCl pH 8.0, 150 mM KCl, 10% glycerol, 1 mM DTT, 0.1 mM ATP, 0.25 mM PMSF) at 4 ℃. The mixture was then incubated with 10 μL 2 mg mL^-1^ TEV protease for 1 h at 4 ℃. The eluted protein was separated from the beads by centrifuging the mixture in Amicon Ultra Free MC tubes at 21,000 *g* for 2 min, flash-frozen in liquid nitrogen, and stored at -80 ℃. A similar procedure was used to purify Lis1, except cells were lysed in a hi-salt phosphate lysis buffer (50 mM potassium phosphate pH = 8.0, 150 mM KAc, 2 mM Mg(Ac)_2,_ 10% glycerol, 10 mM Imidazole, 150 mM NaCl, 50 mM 2-Mercaptoethanol (BME), 1 mM PMSF, 0.2% Triton) and protein was eluted in nu-TEV buffer (50 mM Tris-HCl pH 8.0, 1 mM EGTA, 150 mM KAc, 150 mM NaCl, 2 mM Mg(Ac)_2_, 10% glycerol, 1 mM DTT, 0.5 mM PMSF). The protein concentration was determined by both 280 nm absorbance and the Bradford assay.

Proteins were labeled with fluorescent dyes after resuspending the lysate with beads and before adding TEV protease. 10 nanomoles of a fluorescent dye were well mixed with the protein-bead mixture and incubated for 1 h at 4 ℃. The column was then washed with 100 mL TEV buffer to remove excess dye before eluting the protein from the beads. The labeling percentage was determined by measuring the protein concentration in Bradford assays and the absorbance under 555 nm and 655 nm excitation for TMR/LD555 and Cy5/LD655 dyes, respectively. The probability, *p* of each SNAP-Lis1 monomer labeled with TMR and LD655 dyes derivatized with benzyl guanine was 0.72 and 0.80 per Lis1 monomer, respectively. The probabilities of a SNAP-Lis1 dimer to be labeled with at least one TMR or LD655 dyes (calculated as 2*p* – *p*^2^) were 0.92 and 0.96, respectively. The probability of a dynein dimer being labeled with a dye was 0.8.

Unlike dynein, dynein_Gal_ showed signs of aggregation and slower motility after flash freezing and storage at -80 ℃. To avoid the freezing and thawing cycle, dynein_Gal_ experiments were performed immediately after protein preparation.

### Microtubule Polymerization and Subtilisin Treatment

Tubulin was purified from pig brains in a 1 M PIPES buffer. 60 ng unlabeled, 60 ng biotinylated, and 1 ng fluorescently-labeled tubulin were diluted to 1 mg mL^-1^ in 120 µL BRB80 buffer (80 mM PIPES pH = 6.8, 2 mM MgCl_2_, 1 mM EGTA) supplemented with 1x polymerization mixture (1mM GTP, 10% DMSO) and polymerized at 37 ℃ for 40 min. The mixture was incubated at 37 ℃ for additional 40 min after adding 1 mM taxol. The mixture was centrifuged at 21,000 *g* for 13 min at room temperature and resuspended in 30 µL BRB80 buffer supplemented with 10 mM taxol and 1 mM DTT. Microtubules were stored in the dark at room temperature and used within two weeks.

For subtilisin treatment, 2.5 mg mL^-1^ polymerized microtubules were stored in the dark for 1 day for elongation and then incubated with a different concentration of subtilisin for 2 h at a 37 ℃ bath. Under these conditions, 50 µg mL^-1^ subtilisin cleaved more than 85% of the tubulin tails without substantially affecting dynein motility along these microtubules (not shown). The proteolytic cleavage was stopped with the addition of 2 mM PMSF. The microtubules were centrifuged at 21,000 *g* for 13 min at room temperature and the pellet was resuspended in BRB80 supplemented with 10 mM taxol.

### Single-molecule motility assays

Flow channels were prepared by placing a multichannel parafilm in between a microscope slide and a cover glass functionalized with PEG/PEG-biotin (Microsurfaces Inc.). The channel was incubated with 20 µL of 2 mg mL^-1^ streptavidin for 2 min and excess streptavidin was removed by washing the channel with 60 µL dynein motility buffer (DMB; DLB supplemented with 2% pluronic acid, 1 mM taxol, 1 mM tris(2-carboxyethyl)phosphine (TCEP)). The channel was then incubated with 20 µL of 2 mg mL^-1^ biotinylated microtubules in DMB for 2 min and unbound microtubules were removed by washing the channel with 80 µL DMB. After 3 min, the channel was then washed with 20 µL stepping buffer (DLB supplemented with 0.4% pluronic acid, 1 mM taxol, 1 mM TCEP, 2 mM ATP, 1% gloxy (glucose oxidase and catalase), 0.5% dextrose, and 50 -150 mM KAc). Dynein and Lis1 were diluted in the stepping buffer and incubated on ice for 10 min. 20 µL motor+Lis1 final mixture in stepping buffer was added to the channel and the sample was imaged immediately for 15 min.

Lis1-dynein colocalization experiments were performed by mixing fluorescently-labeled dynein and Lis1 into the final mixture, before flowing into the chamber. Colocalization between two differentially labeled dynein was performed by mixing 5 nM LD555 dynein with 5 nM LD655-dynein and letting on ice for 10 min with or without Lis1. To maximize crosslinking efficiency with different colors of dyneins, dyneins were added to the mixture before Lis1. The dynein-Lis1 mixture was diluted in stepping buffer supplemented with 1 mM ATP and 50 mM KAc.

Single-molecule imaging was performed using a custom-built objective-type TIRF microscope equipped with an inverted microscopy body (Nikon Ti-Eclipse), 100× magnification 1.49 numerical aperture (N.A.) planapochromat oil-immersion objective (Nikon), and a perfect focusing system. The fluorescence signal was detected using an electron-multiplied charge-coupled device (EM-CCD) camera (Andor, Ixon). The effective pixel size after magnification was 108 nm. Samples labeled with GFP, TMR/LD555, and Cy5/LD655 were excited using 0.04 kW cm^-2^ 488, 532, and 633 nm laser beams (Coherent), and the emission signal was detected using bandpass emission filters (Semrock). For two- and three-color fluorescence assays, imaging was performed using alternating excitation and the time-sharing mode.

Single-color movies were recorded at 1 s per frame, whereas multi-color movies were recorded with 0.5 s per frame per color using the time-sharing mode. To introduce or remove Lis1 from the channel during imaging, two 0.5 mm diameter holes were drilled on a glass slide at both ends of the flow channel. The region of interest in the channel was imaged for 1 min before introducing Lis1 or washing the free Lis1 from the channel while imaging dynein motility in real-time.

### Data Analysis

Movies were analyzed in ImageJ to create kymographs. Motors that are stationary, exhibit diffusional movement, or move less than 3 pixels were excluded from the analysis. The results of the data analysis were plotted in Prism and Origin. Data fitting was performed in Origin. Cumulative distribution functions (CDFs) of the run length and run-time data were fitted to a double exponential decay, *y* = *A*_l_*e*^-*t*/*τ*_l_^ + *A*_2_*e*^-*t*/τ_2_^ where *A_l_* and *A_2_* are the amplitudes and *τ_l_* and *τ_2_* are the decay constants. The weighted average of the decay constants, 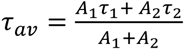 was reported as the decay constant^55^.

The average velocities of motors that colocalize with 0 (*V_O_*), 1 (*V_l_*), and 2 (*V_2_*) colors of Lis1 were determined from three-color TIRF assays. To determine *K_D_* of Lis1, the average dynein velocity under different Lis1 concentrations, *V ([Lis*1]) was fitted to (1 *- p_b_)^2^V_O_ +* 2*p_b_(*1 *- p_b_)V_l_ + (p_b_)^2^V_2_* where 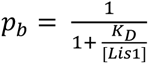 is the binding probability of Lis1 to a dynein monomer. The probability of Lis1 colocalization to dynein_Gal_ in Extended Data Figure 4c was fitted to a binding isotherm function, defined as *C(*2*p_b_ - (p_b_)^2^)*, where *C* is less than 1 due to incomplete labeling of Lis1 and the limited single-molecule detection ability due to the presence of labeled Lis1 in the solution.

### Microtubule co-pelleting assays

Unlabeled taxol-stabilized microtubules were polymerized as above and free tubulin was removed by centrifugation through a 60% glycerol cushion in BRB80 (80 mM PIPES-KOH pH 6.8, 1 mM MgCl_2_, 1 mM EGTA, 1 mM DTT, 20 µM taxol) for 15 min at 100,000xg and 37 ℃. The microtubule pellet was resuspended in DLB supplemented with 20 µM taxol. Microtubules (0 - 600nM tubulin) were incubated with 100 nM Lis1 for 10 min before being pelleted for 15 min at 100,000 *g* and 25 ℃. The supernatant was analyzed via SDS-PAGE and depletion was determined using densitometry in ImageJ. Binding curves were fit in binding isotherm in Origin.

### Optical trapping assays

Optical trapping experiments were performed on a custom-built optical trap using a Nikon Ti-E microscope body and a 100× 1.49 NA planapochromat oil immersion objective, as previously described^31^. The beads were trapped with a 2W 1,064 nm laser (IPG Photonics) and the trap was steered with a two-axis acousto-optical deflector (AA Electronics). The trap stiffness was calculated from the Lorentzian fit to the power spectrum of a trapped bead. For stall force measurements, the trap stiffness was adjusted to allow the motor to walk 100 nm away from the trap center on average before the bead comes to the stall. The position of the bead from the center of the fixed trap was recorded for at least 90 s at 5 kHz.

For fixed trap measurements, 0.8 µm diameter carboxylated latex beads (Life Technologies) were functionalized with rabbit polyclonal GFP antibody (BioLegend, no. MMS-118P), as previously described^31^. In optical trapping assays, 2 mM and 0.25 mM casein was used instead of 2% and 0.4% pluronic acid in DMB and stepping buffer, respectively. The 2 μL of the bead stock were diluted in 8 µL DLB. 2 µL of diluted beads were sonicated for 8 s. To ensure that more than 90% of the beads are driven by single motors, GFP-dynein concentration was diluted in the stepping buffer to desired concentrations such that less than 30% of the beads exhibit motility when brought on top of a surface-immobilized axoneme. The mixture was sonicated for 8 s and 2 µL of diluted beads were mixed with 2 µL GFP-dynein, 1.5 µL Lis1, and 2 µL stepping buffer, and incubated on ice for 10 min. For the no Lis1 condition, an equal volume of stepping buffer was added to the mixture instead of Lis1.

Cy5-labeled sea urchin axonemes were nonspecifically adsorbed to the channel surface. After 30 s incubation, the channel was washed with 60 μL DMB to remove the unbound axoneme. After 3 min incubation, the channel was washed with 60 μL DMB and then with 20 μL stepping buffer. 7.5 μL of the dynein-Lis1-bead mixture was diluted in 20 µL stepping buffer and the KAc concentration was adjusted to 50 mM before being flown into the channel. The channel was sealed with nail polish to prevent evaporation during data acquisition.

Optical trap data were analyzed with a custom-written MATLAB script. The data were downsampled to 500 Hz via median filtering before analysis. Stall events were manually scored when the velocity of the bead movement was reduced to ∼0 nm s^-1^ under hindering force and terminated with a sudden (<4 ms) snapping of the bead to the trap center. Events that occur within a 25 nm distance to the trap center, last shorter than 0.4 s, or terminate with backward movement, multiple-step detachment, or slow (>4 ms) return of the bead to the trap center were excluded from the analysis. Stall forces were calculated as the average force value of the plateau where the bead is nearly immobile before detaching from the axoneme. CDFs of stall times were fitted to a double exponential decay and the weighted average of two decay constants from the fit was reported as the decay constant.

### F-V measurements

Similar sample preparation procedures were used for F-V measurements using an optical trap. After the bead movement reached half of the average stall force of the motor, the trapping beam repositioned itself and maintained a 100 nm distance from the bead via a force-feedback mechanism. To determine the microtubule polarity in the no-ATP condition, LD655-labeled GST-Dyn_331kDA_ motors (without a GFP tag) were flown into the channel in a stepping buffer supplemented with 30 μM ATP. The motors were allowed to walk on and accumulate at the minus-end of surface-immobilized axonemes for 4 mins. The channel was then washed with 40 µL DLBM buffer, followed by 20 µL stepping buffer supplemented with 0.5 U mL^-1^ apyrase instead of ATP to deplete residual ATP in the channel. 0.5 U mL^-1^ apyrase was also included in the bead-dynein-Lis1 mixture to remove residual ATP from protein preparations. During optical trapping, microtubule polarity was determined by the minus-end accumulation of the LD655 signal detected by an sCMOS camera (Hamamatsu, Orca Flash 4.0). Beads were pulled along the length of an axoneme at a constant velocity, and the beads that engaged with the axoneme were trapped by repositioning the trapping beam 100 nm away from the bead center along the direction of the applied force.

The bead position was recorded at 5 kHz and the data were downsampled to 500 Hz via median filtering. The slope of each trace was defined as the velocity of individual motors under an applied force. The traces that are shorter than 70 ms, show bidirectional motility under force, or exhibit instantaneous jumps larger than 50 nm were excluded from data analysis. The asymmetry ratios were calculated by comparing the average velocities under forces equally larger or smaller than *F_stall_* in the presence of ATP, and forces equally larger or smaller than 0 pN in the absence of ATP. Velocities at corresponding forces were either directly measured or calculated by linear regression of the measured velocities under the closest force measurements. The errors of asymmetry ratios were determined by error propagation of the compared velocities.

### Statistical Analysis

The p-values were calculated by the two-tailed t-test for stall force histograms, the two-tailed t-test with Welch corrections for velocity measurements, and the Kolmogorov-Smirnov test for run length and run time measurements in Prism and Origin. CDFs were calculated in MATLAB.

### Data Availability

A reporting summary for this article is available as Supplementary Information file. The main data supporting the findings of this study are available within the article and its Supplementary Figures. The source data and protocols that support the findings of this study will be made available by the corresponding authors upon reasonable request.

## Extended Data

**Extended Data Table 1.**
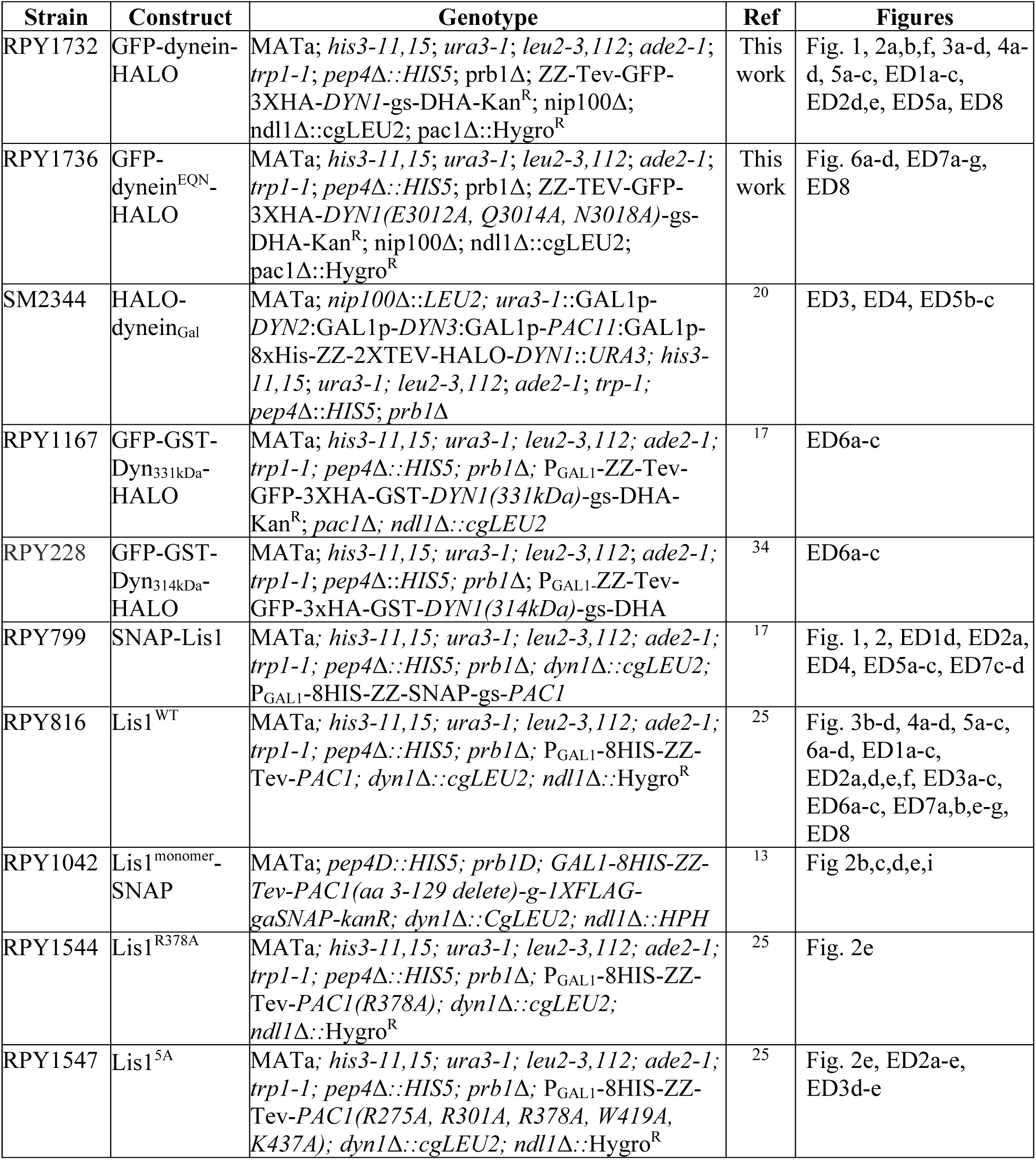
The list of *S. cerevisiae* strains used in this study. SNAP and HALO (DHA) are genetic tags used for labeling the protein with a fluorescent dye. Lis1 constructs that are not tagged with SNAP were labeled with maleimide reactive probes. TEV indicates a TEV protease cleavage site. P_GAL1_ denotes the galactose promoter, which was used for inducing strong expression of Lis1 and dynein motor domain constructs. Amino acid spacers are indicated by g (glycine) and gs (glycine-serine). (Ref: Reference, ED: Extended Data Figure).

## Extended Data Figures

**Extended Data Figure 1.**
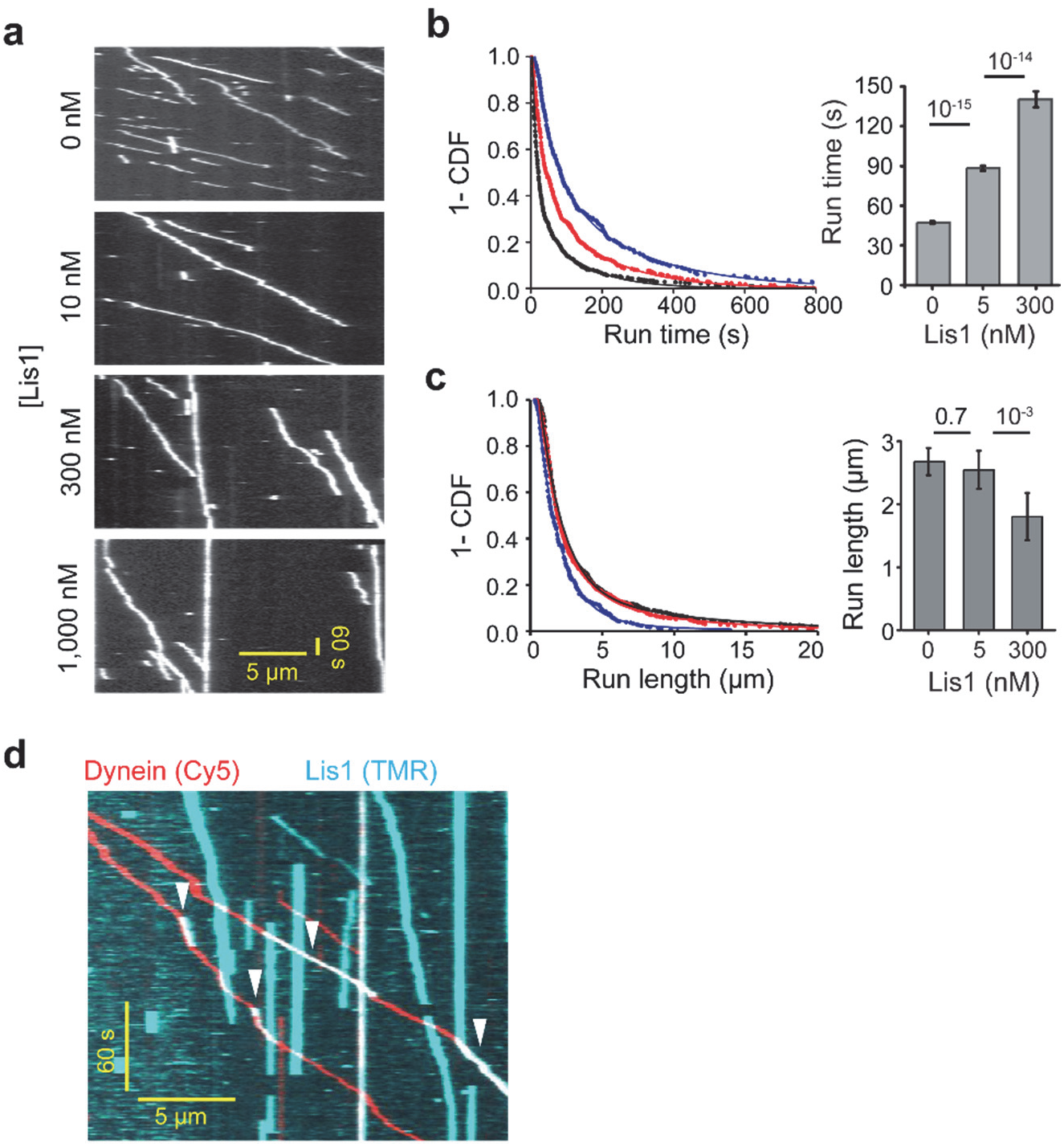
Single molecule motility of dynein in the presence of Lis1. **a)** Representative kymographs of TMR-dynein with increasing concentrations of unlabeled Lis1. **b)** (Left) 1-CDF of run time under different Lis1 concentrations. Fitting to a double exponential decay (solid curves) reveals the weighted average of run time in each condition. (Right) The weighted average of run time under increasing Lis1 concentrations (± s.e.; N = 861, 534, and 312 from left to right). **c)** (Left) 1-CDF of run length under different Lis1 concentrations. Solid curves represent a fit to a double exponential decay. (Right) The weighted average of run length in each condition (bar graphs, ± s.e.; N = 861, 534, and 312 from left to right). **d)** An example kymograph shows that the transient binding of Lis1 slows down whereas the subsequent release of Lis1 restores the velocity (arrowheads). In **b** and **c**, P values were calculated by the Kolmogorov-Smirnov test.

**Extended Data Figure 2.**
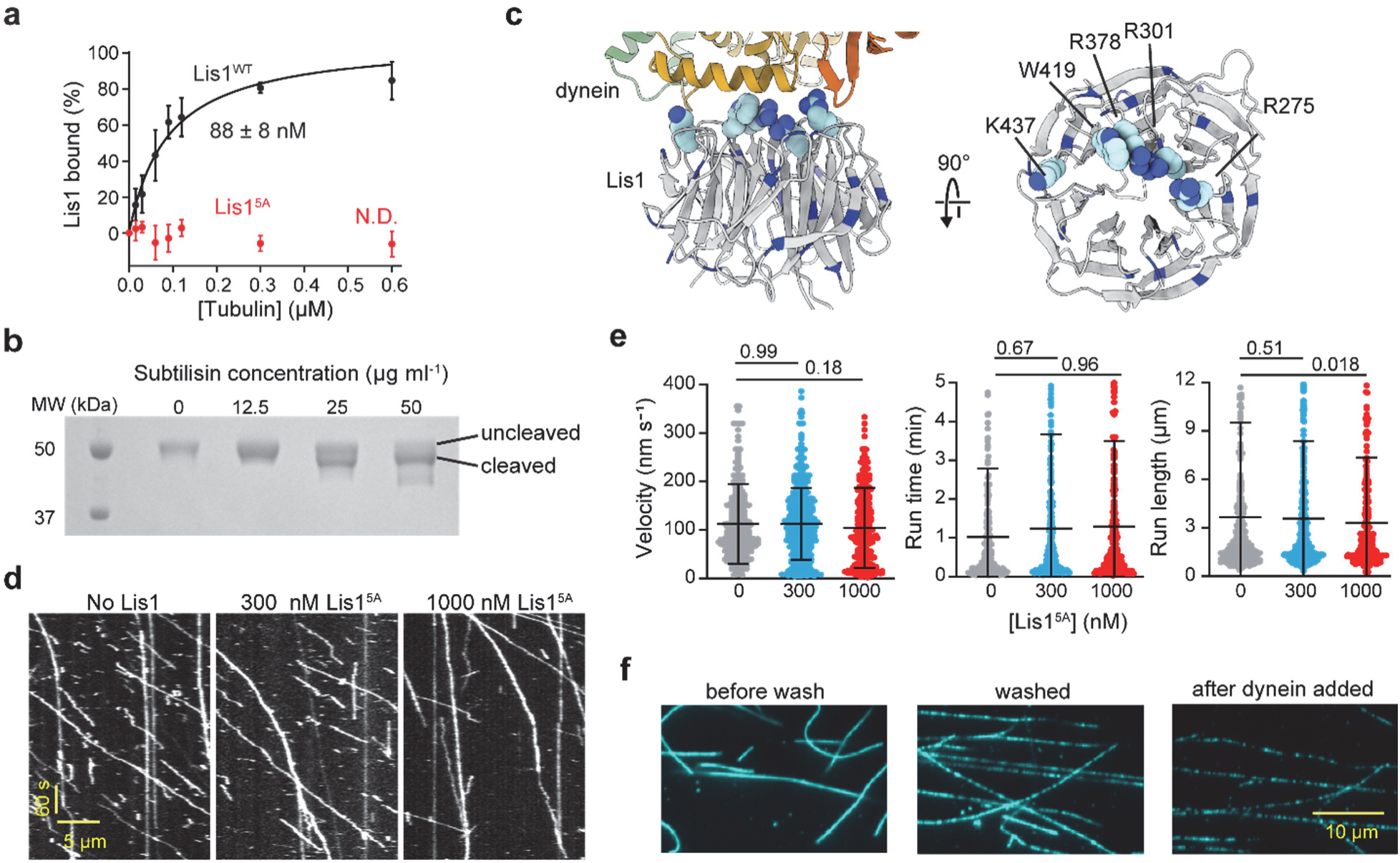
Lis1 interacts with the microtubule lattice through its dynein binding site in vitro. **a)** Microtubule co-pelleting assay with Lis1^WT^ and Lis1^5A^ (mean ±s.e.m., three replicates per condition). The solid curve represents a fit to a binding isotherm to determine K_D_ (±s.e.; N.D.: not determined). The statistical analysis was performed using an extra sum-of-squares F test (p <0.0001). **b**) The subtilisin treatment of microtubules reduces the molecular weight of tubulin in a denaturing gel. **c)** The structure of Lis1 bound to the AAA ring with the five residues mutated in Lis1^5A^ are shown as spheres and all lysine and arginine residues are in blue. **d)** Representative kymographs of dynein in the presence and absence of Lis1^5A^. Assays were performed in 50 mM KAc. **e)** The velocity, run time, and run length of dynein in the presence and absence of Lis1^5A^ (N = 326, 534, and 336 from left to right). The center line and whiskers represent the mean and s.d., respectively. P values were calculated by a two-tailed t-test with Welch correction for velocity and by Kolmogorov-Smirnov test for run time and run length. **f)** Surface-immobilized microtubules were decorated by 100 nM TMR-Lis1 before and after removing unbound Lis1 in the channel. The assay was performed in the absence of added salt to maximize the Lis1-microtubule interaction. Washing the chamber with buffer and introducing dynein reduces 92% of the Lis1 signal on microtubules.

**Extended Data Figure 3.**
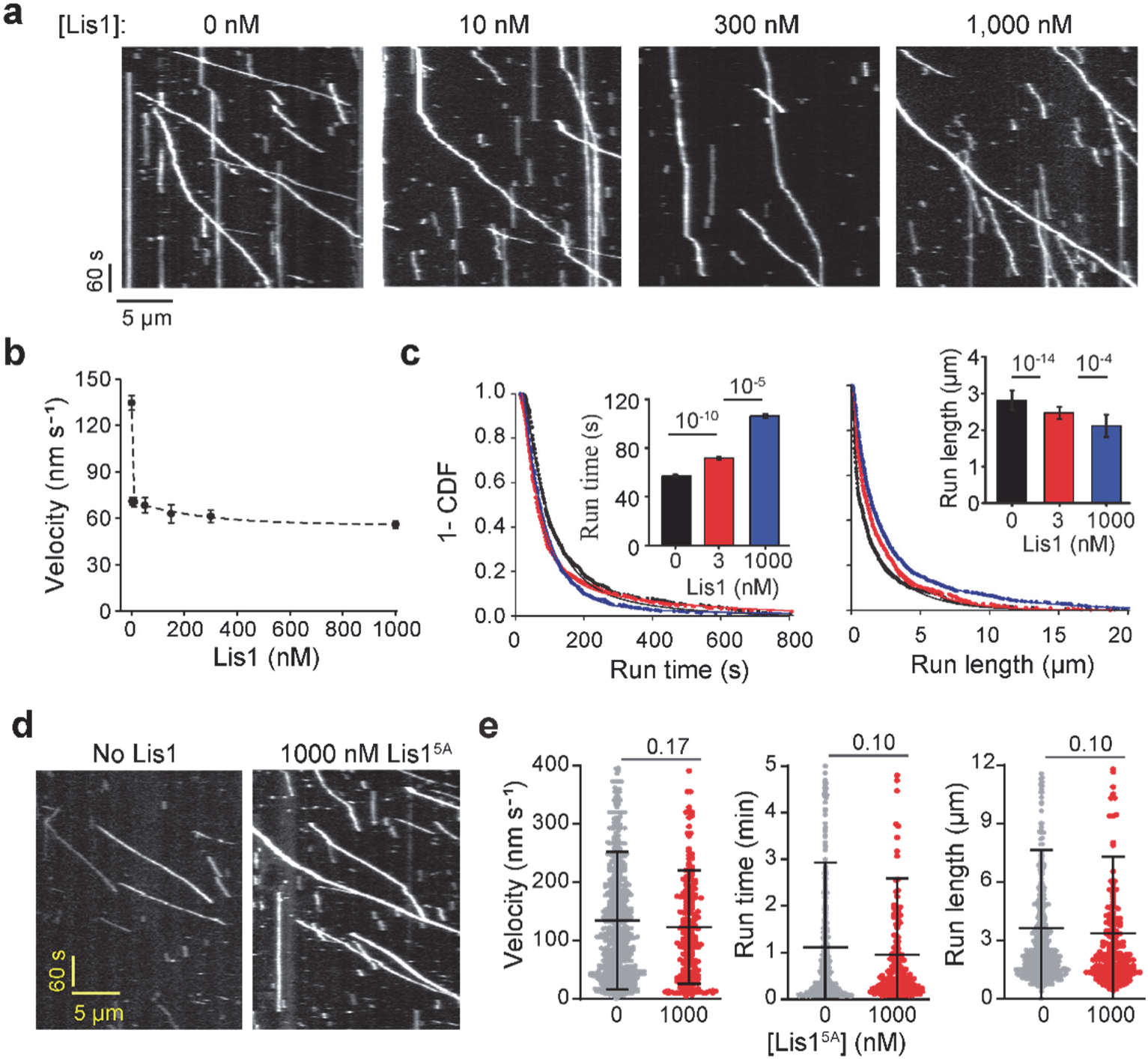
Motility of dynein_Gal_ in the presence of unlabeled Lis1. **a)** Representative kymographs of dynein_Gal_ with increasing concentrations of unlabeled Lis1. Assays were performed in 1 mM ATP and 150 mM KAc. **b)** The velocity of dynein_Gal_ under different Lis1 concentrations (mean ± s.e.m.; N = 611, 694, 230, 260, 251, and 852 from left to right; two biological replicates). **c)** 1-CDF of run time and run length of dynein_Gal_ under different Lis1 concentrations. Fitting to a double exponential decay (solid curves) reveals the weighted average of run time and run length of the motor in each condition (bar graphs, ± s.e.; N = 611, 694, and 852 from left to right). **d)** Representative kymographs of dynein_Gal_ in the presence or absence of 1,000 nM unlabeled Lis1^5A^. **e**) The velocity, run time, and run length of dynein_Gal_ in the presence or absence of 1,000 nM Lis1^5A^ (N = 610 and 209 from left to right). The center line and whiskers represent the mean and s.d., respectively. In **c** and **e**, P values were calculated by a two-tailed t-test with Welch correction for velocity and by Kolmogorov-Smirnov test for run time and run length.

**Extended Data Figure 4.**
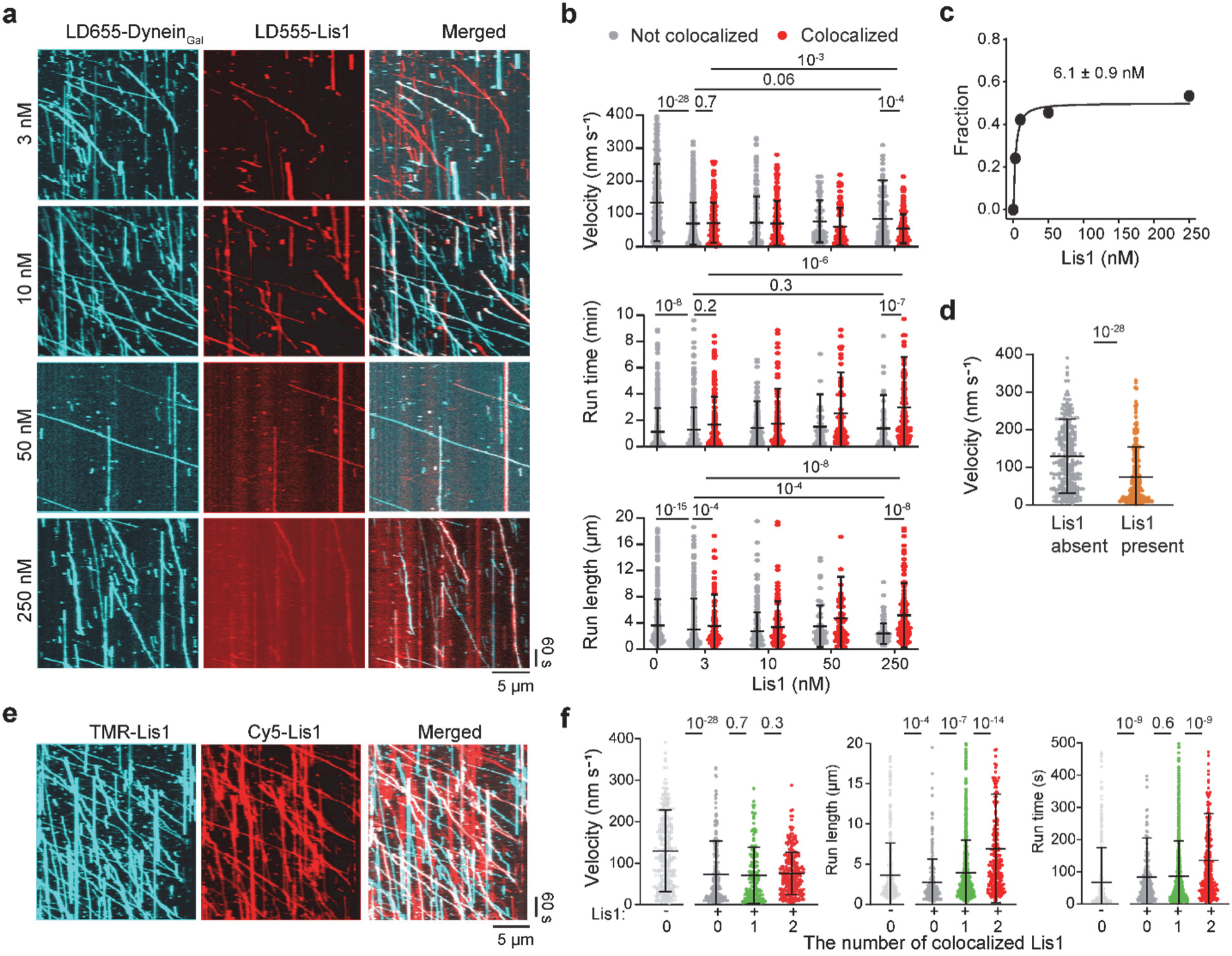
Motility of dynein_Gal_ in the presence of fluorescently-labeled Lis1. **a)** Representative kymographs of two-color imaging of LD655-labeled dynein_Gal_ and LD555-labeled Lis1 under different Lis1 concentrations. **b)** The velocity, run time, and run length distributions of dynein_Gal_ under different concentrations of Lis1 (N = 611, 526, 168, 233, 175, 55, 69, 126, and 145 from left to right). **c)** The fraction of LD655-dynein_Gal_ that colocalizes with LD555-Lis1 under different Lis1 concentrations. A fit to a binding isotherm function (solid curve, see Methods) reveals K_D_ (±s.e.). **d)** The velocity distribution of dynein_Gal_ in the absence of Lis1 compared to the motors that do not colocalize with Lis1 when fluorescently labeled Lis1 is present in the chamber (N = 611 and 233 from left to right). **e)** Representative kymographs show colocalization of 5 nM TMR-labeled and 5 nM Cy5-labeled Lis1 to unlabeled dynein_Gal_. **f)** Velocity, run length, and run time distributions of dynein_Gal_ in the absence and presence of Lis1 in the channel (N = 287, 233, 175, and 274 from left to right). Dynein_Gal_ motors that colocalize with 0, 1, or 2 colors of Lis1 were analyzed separately. In **b**, **d**, and **f**, the center line and whiskers represent the mean and s.d., respectively. P values were calculated by two-tailed t-tests with Welch correction for velocity and by Kolmogorov-Smirnov test for run time and run-length measurements.

**Extended Data Figure 5.**
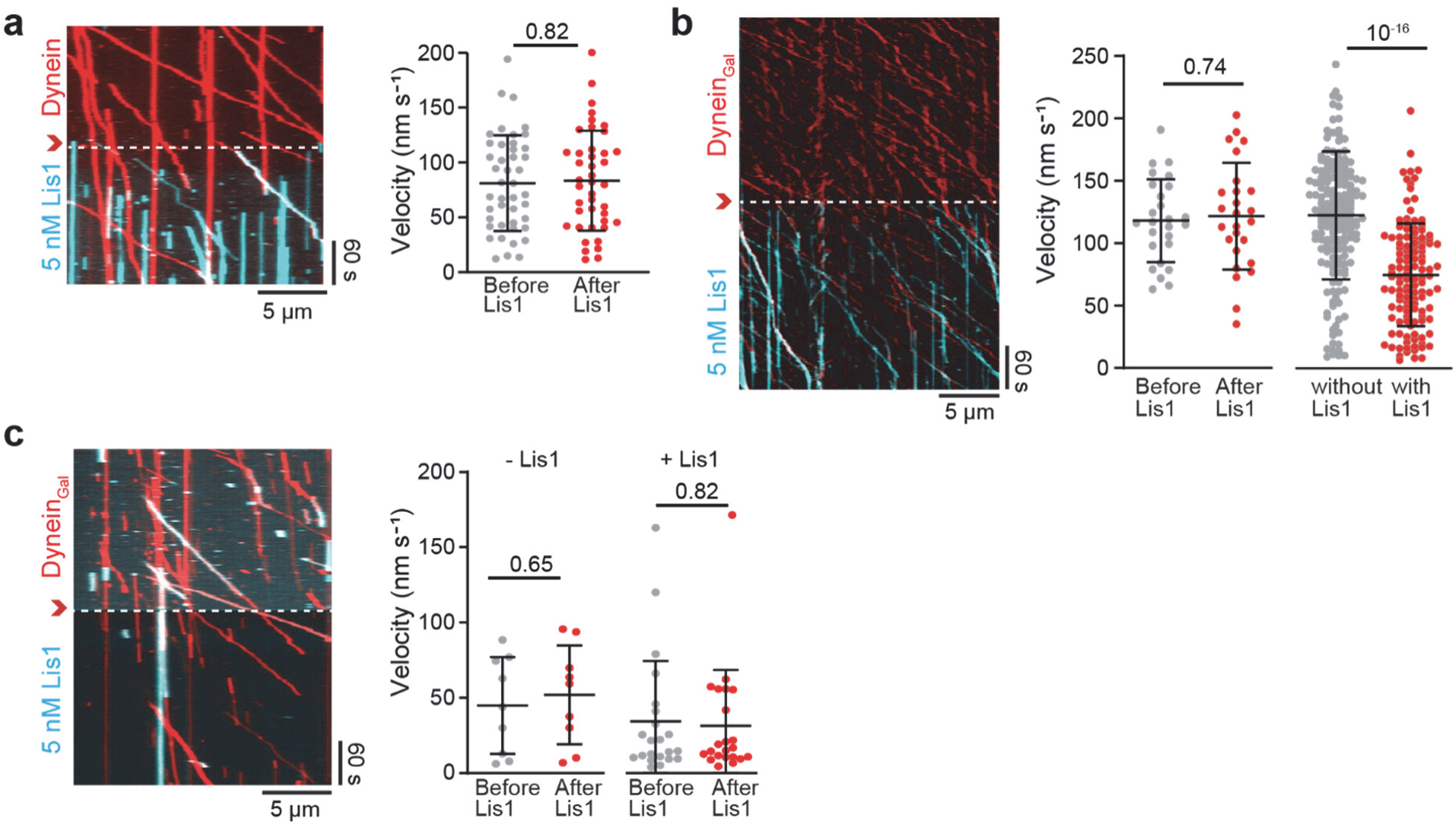
The addition and removal of Lis1 whilst recording dynein motility. **a)** A kymograph (left) and velocity (right) of dynein before and after Lis1 was flown into the flow chamber (red arrowhead and a dashed line, N = 44 and 42 from left to right). **b)** A kymograph of dynein_Gal_ when Lis1 was flown into the channel (red arrowhead and a dashed line). (Middle) The velocity of the complexes that were already walking on the microtubules during Lis1 addition moved at the same velocity after flowing Lis1 (N = 26 and 26 from left to right). Only the motors that do not colocalize with Lis1 were included in the analysis. (Right) The velocity of the complexes that land onto microtubules within 4 minutes after Lis1 addition was analyzed (N = 122 and 84 from left to right). The motors that colocalize with Lis1 walked slower than motors that do not colocalize with Lis1. **c)** A kymograph (left) and velocity (right) of dynein_Gal_ before and after washing excess Lis1 from the flow chamber (red arrowhead and a dashed line). The velocity of the motors that colocalize (+Lis1) or do not colocalize (-Lis1) with Lis1 remained unaltered by removing excess Lis1 from the chamber (N = 9, 9, 22, 22 from left to right). In **a**, **b**, and **c**, assays were performed in 1 mM ATP and 50 mM KAc. The center line and whiskers represent the mean and s.d., respectively. P-values were calculated by a two-tailed t-test with Welch correction.

**Extended Data Figure 6.**
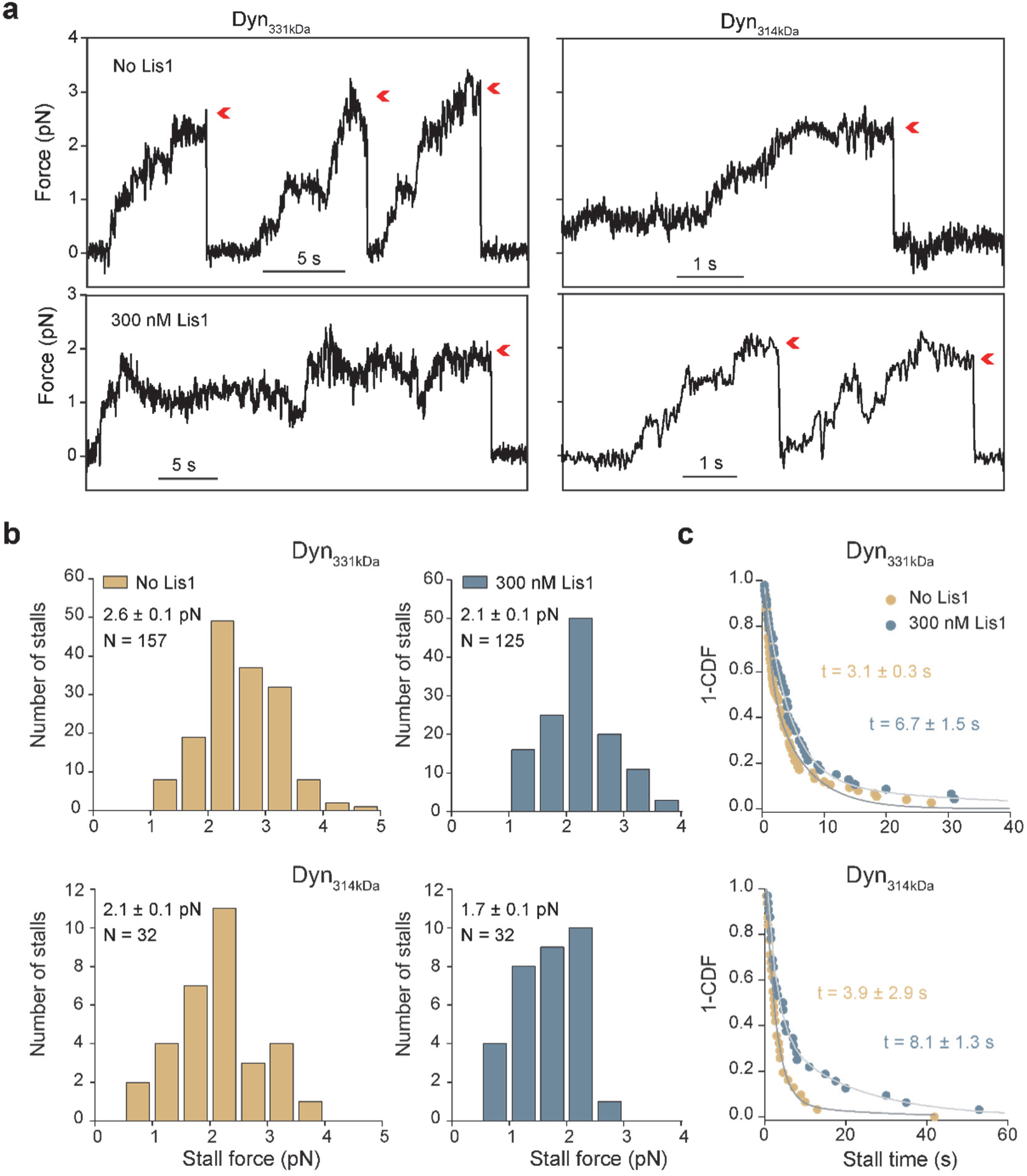
Lis1 reduces the stall force of the tail-truncated dynein. **a)** Representative trajectories of beads driven by tail-truncated dynein constructs dimerized with a Glutathione S-transferase (GST) tag (GFP-GST-Dyn_331kDa_ and GFP-GST-Dyn_314kDa_) in the presence and absence of 300 nM Lis1. Assays were performed in 2 mM ATP. Red arrowheads represent the detachment of the motor from the microtubule followed by the snapping back of the bead to the trap center. **b)** Stall force histograms of Dyn_331kDa_ and Dyn_314kDa_ in the presence and absence of 300 nM Lis1 (mean ± s.e.m.; p = 2×10^-7^ for Dyn_331kDa_ and 0.006 for Dyn_314kDa_, two-tailed t-test). **c)** Stall times of Dyn_331kDa_ and Dyn_314kDa_ in the presence and absence of 300 nM Lis1. Fitting to a double exponential decay (solid curves) reveals the weighted average of stall time (±s.e.).

**Extended Data Figure 7.**
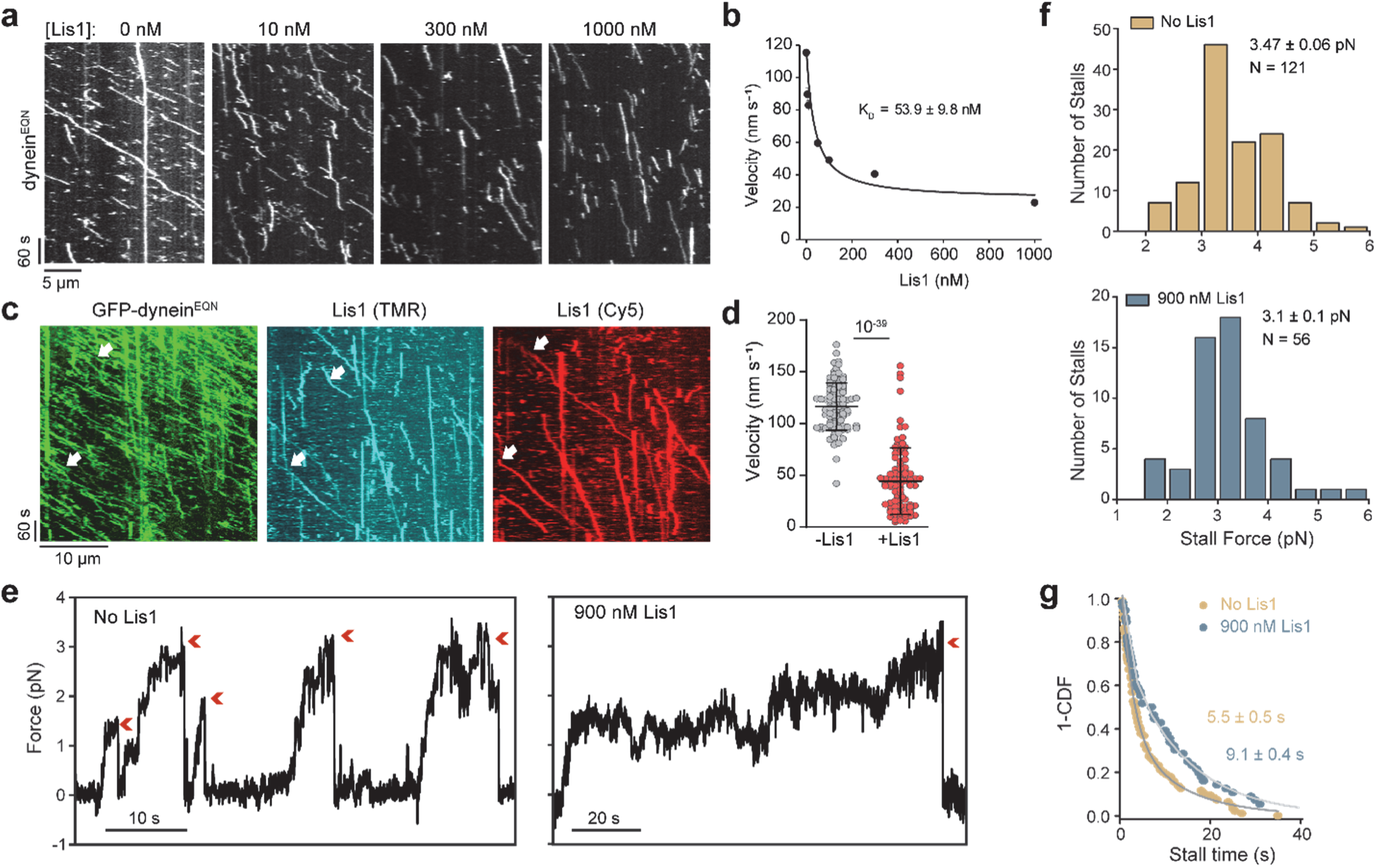
Lis1 slows the motility but does not substantially affect the stall force of dynein^EQN^. **a)** Representative kymographs of dynein^EQN^ motility under different concentrations of unlabeled Lis1. **b)** The velocity of dynein^EQN^ motility under different concentrations of unlabeled Lis1 (mean ± s.e.m.; N = 659, 283, 482, 484, 487, 610, and 132 from left to right). The fit of the dynein^EQN^ velocity data (solid curve) reveals K_D_ (±s.e., see Methods). **c)** A representative kymograph of a three-color imaging assay shows two Lis1s bind to the same dynein^EQN^ motor (white arrows). **d)** The velocities of dynein^EQN^ motors not colocalizing (-Lis1) or colocalizing (+Lis1) with Lis1 (N = 98 and 92 from left to right). The center line and whiskers represent the mean and s.d., respectively. The P-value was calculated by a two-tailed t-test with Welch correction. **e)** Representative trajectories of beads driven by dynein^EQN^ in the presence or absence of 900 nM Lis1 in a fixed trapping assay. Assays were performed in 2 mM ATP. Red arrowheads represent the detachment of the motor from the microtubule followed by the snapping back of the bead to the trap center. **f)** Stall force histograms of dynein^EQN^ in the presence and absence of 900 nM Lis1 (mean ± s.e.m.; p = 10^-3^, two-tailed t-test). **g)** Stall times of dynein^EQN^ in the presence and absence of 900 nM Lis1. Fitting to a double exponential decay (solid curves) reveals the weighted average of stall time (±s.e.).

**Extended Data Figure 8.**
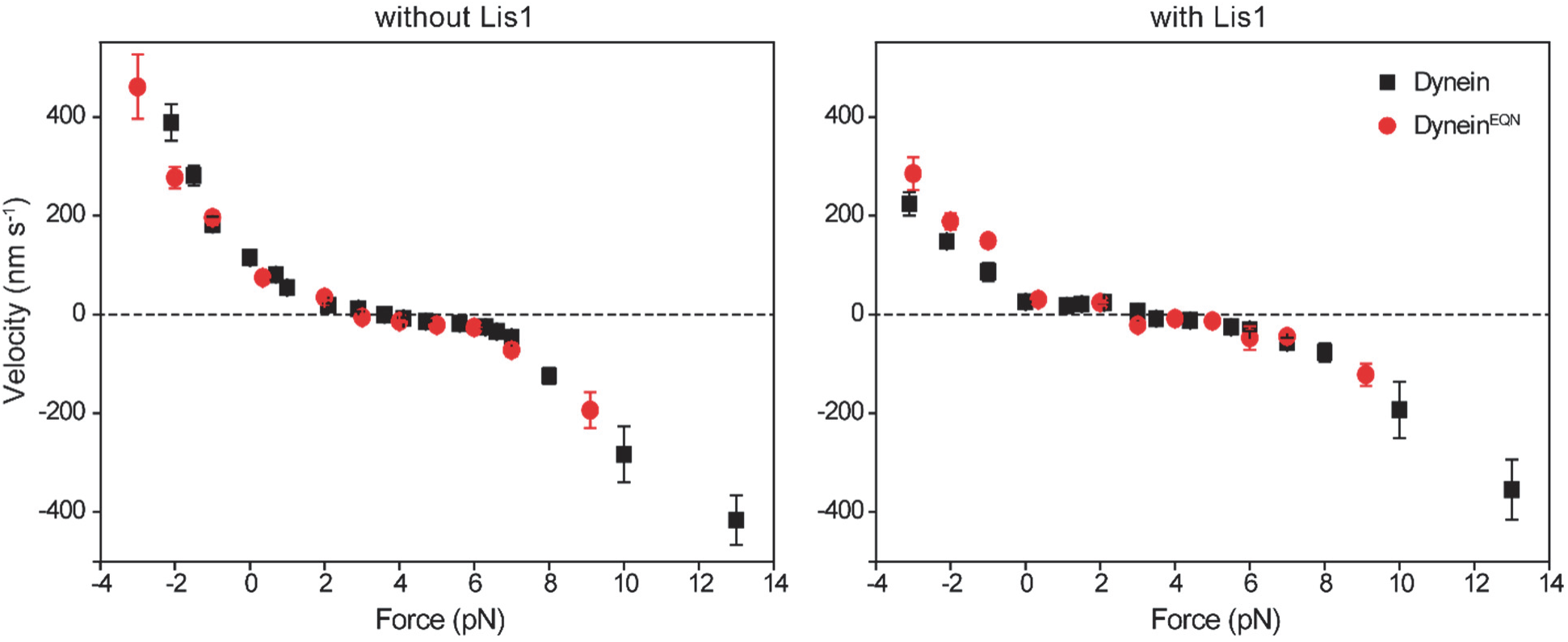
The comparison of the F-V behavior of dynein and dynein^EQN^ in the presence and absence of Lis1. (Left) F-V measurements of dynein and dynein^EQN^ in the absence of Lis1 (mean ±s.e.m., from left to right, N = 41, 38, 37, 861, 27, 52, 106, 28, 51, 60, 89, 115, 124, 62, 23, 38, 40 for dynein and N = 31, 66, 62, 48, 47, 26, 42, 39, 23, 45, 33 for dynein^EQN^). (Right) F-V measurements of dynein in 300 nM Lis1 compared to F-V of dynein^EQN^ in 900 nM Lis1 (mean ±s.e.m., from left to right, N = 34, 52, 25, 312, 33, 42, 49, 37 62, 47 33 42, 36, 29, 18 for dynein; and N = 54, 36, 61, 31, 37, 17, 54, 14, 19, 38, 26 for dynein^EQN^). Dynein velocity under assisting forces is lower than that of dynein^EQN^ in the presence of Lis1. Assays were performed in 2 mM ATP.

## References

1. Canty, J.T., Tan, R., Kusakci, E., Fernandes, J. & Yildiz, A. Structure and Mechanics of Dynein Motors. Annual review of biophysics 50, 549–574 (2021).

2. Reck-Peterson, S.L., Redwine, W.B., Vale, R.D. & Carter, A.P. The cytoplasmic dynein transport machinery and its many cargoes. Nature reviews. Molecular cell biology 19, 382–398 (2018).

3. Guedes-Dias, P. & Holzbaur, E.L.F. Axonal transport: Driving synaptic function. Science 366, eaaw9997 (2019).

4. Carter, A.P., Cho, C., Jin, L. & Vale, R.D. Crystal structure of the dynein motor domain. Science 331, 1159–1165 (2011).

5. Kon, T. et al. The 2.8 A crystal structure of the dynein motor domain. Nature 484, 345–350 (2012).

6. Can, S., Lacey, S., Gur, M., Carter, A.P. & Yildiz, A. Directionality of dynein is controlled by the angle and length of its stalk. Nature 566, 407–410 (2019).

7. Kon, T. et al. Helix sliding in the stalk coiled coil of dynein couples ATPase and microtubule binding. Nature structural & molecular biology 16, 325–333 (2009).

8. Torisawa, T. et al. Autoinhibition and cooperative activation mechanisms of cytoplasmic dynein. Nature cell biology 16, 1118–1124 (2014).

9. Trokter, M., Mucke, N. & Surrey, T. Reconstitution of the human cytoplasmic dynein complex. Proceedings of the National Academy of Sciences of the United States of America 109, 20895–20900 (2012).

10. Zhang, K. et al. Cryo-EM Reveals How Human Cytoplasmic Dynein Is Auto-inhibited and Activated. Cell 169, 1303–1314 e1318 (2017).

11. McKenney, R.J., Huynh, W., Tanenbaum, M.E., Bhabha, G. & Vale, R.D. Activation of cytoplasmic dynein motility by dynactin-cargo adapter complexes. Science 345, 337–341 (2014).

12. Schlager, M.A., Hoang, H.T., Urnavicius, L., Bullock, S.L. & Carter, A.P. In vitro reconstitution of a highly processive recombinant human dynein complex. The EMBO journal 33, 1855–1868 (2014).

13. Huang, J., Roberts, A.J., Leschziner, A.E. & Reck-Peterson, S.L. Lis1 acts as a "clutch" between the ATPase and microtubule-binding domains of the dynein motor. Cell 150, 975–986 (2012).

14. Reiner, O. et al. Isolation of a Miller-Dieker lissencephaly gene containing G protein beta-subunit-like repeats. Nature 364, 717–721 (1993).

15. Markus, S.M., Marzo, M.G. & McKenney, R.J. New insights into the mechanism of dynein motor regulation by lissencephaly-1. eLife 9, 59737 (2020).

16. Gillies, J.P. et al. Structural basis for cytoplasmic dynein-1 regulation by Lis1. eLife 11, 71229 (2022).

17. DeSantis, M.E. et al. Lis1 Has Two Opposing Modes of Regulating Cytoplasmic Dynein. Cell 170, 1197–1208 e1112 (2017).

18. Elshenawy, M.M. et al. Lis1 activates dynein motility by modulating its pairing with dynactin. Nature cell biology 22, 570–578 (2020).

19. Htet, Z.M. et al. LIS1 promotes the formation of activated cytoplasmic dynein-1 complexes. Nature cell biology 22, 518–525 (2020).

20. Marzo, M.G., Griswold, J.M. & Markus, S.M. Pac1/LIS1 stabilizes an uninhibited conformation of dynein to coordinate its localization and activity. Nature cell biology 22, 559–569 (2020).

21. Qiu, R., Zhang, J. & Xiang, X. LIS1 regulates cargo-adapter-mediated activation of dynein by overcoming its autoinhibition in vivo. J Cell Biol 218, 3630–3646 (2019).

22. Yamada, M. et al. LIS1 and NDEL1 coordinate the plus-end-directed transport of cytoplasmic dynein. The EMBO journal 27, 2471–2483 (2008).

23. Torisawa, T. et al. Functional dissection of LIS1 and NDEL1 towards understanding the molecular mechanisms of cytoplasmic dynein regulation. The Journal of biological chemistry 286, 1959–1965 (2011).

24. Wang, S. et al. Nudel/NudE and Lis1 promote dynein and dynactin interaction in the context of spindle morphogenesis. Molecular biology of the cell 24, 3522–3533 (2013).

25. Toropova, K. et al. Lis1 regulates dynein by sterically blocking its mechanochemical cycle. eLife 3, 03372 (2014).

26. Baumbach, J. et al. Lissencephaly-1 is a context-dependent regulator of the human dynein complex. eLife 6, 21768 (2017).

27. Gutierrez, P.A., Ackermann, B.E., Vershinin, M. & McKenney, R.J. Differential effects of the dynein-regulatory factor Lissencephaly-1 on processive dynein-dynactin motility. The Journal of biological chemistry 292, 12245–12255 (2017).

28. DeWitt, M.A., Chang, A.Y., Combs, P.A. & Yildiz, A. Cytoplasmic dynein moves through uncoordinated stepping of the AAA+ ring domains. Science 335, 221–225 (2012).

29. Qiu, W. et al. Dynein achieves processive motion using both stochastic and coordinated stepping. Nature structural & molecular biology 19, 193–200 (2012).

30. Belyy, V. et al. The mammalian dynein-dynactin complex is a strong opponent to kinesin in a tug-of-war competition. Nature cell biology 18, 1018–1024 (2016).

31. Belyy, V., Hendel, N.L., Chien, A. & Yildiz, A. Cytoplasmic dynein transports cargos via load-sharing between the heads. Nature communications 5, 5544 (2014).

32. Muhua, L., Karpova, T.S. & Cooper, J.A. A yeast actin-related protein homologous to that in vertebrate dynactin complex is important for spindle orientation and nuclear migration. Cell 78, 669–679 (1994).

33. Heil-Chapdelaine, R.A., Oberle, J.R. & Cooper, J.A. The cortical protein Num1p is essential for dynein-dependent interactions of microtubules with the cortex. J Cell Biol 151, 1337–1344 (2000).

34. Reck-Peterson, S.L. et al. Single-molecule analysis of dynein processivity and stepping behavior. Cell 126, 335–348 (2006).

35. McKenney, R.J., Vershinin, M., Kunwar, A., Vallee, R.B. & Gross, S.P. LIS1 and NudE induce a persistent dynein force-producing state. Cell 141, 304–314 (2010).

36. Markus, S.M. & Lee, W.L. Regulated offloading of cytoplasmic dynein from microtubule plus ends to the cortex. Developmental cell 20, 639–651 (2011).

37. Monroy, B.Y. et al. A Combinatorial MAP Code Dictates Polarized Microtubule Transport. Developmental cell 53, 60–72 e64 (2020).

38. Ferro, L.S. et al. Structural and functional insight into regulation of kinesin-1 by microtubule-associated protein MAP7. Science 375, 326–331 (2022).

39. Gennerich, A., Carter, A.P., Reck-Peterson, S.L. & Vale, R.D. Force-induced bidirectional stepping of cytoplasmic dynein. Cell 131, 952–965 (2007).

40. Rao, L., Berger, F., Nicholas, M.P. & Gennerich, A. Molecular mechanism of cytoplasmic dynein tension sensing. Nature communications 10, 3332 (2019).

41. Cleary, F.B. et al. Tension on the linker gates the ATP-dependent release of dynein from microtubules. Nature communications 5, 4587 (2014).

42. Nicholas, M.P. et al. Cytoplasmic dynein regulates its attachment to microtubules via nucleotide state-switched mechanosensing at multiple AAA domains. Proceedings of the National Academy of Sciences of the United States of America 112, 6371–6376 (2015).

43. Ezber, Y., Belyy, V., Can, S. & Yildiz, A. Dynein harnesses active fluctuations of microtubules for faster movement. Nature Physics 16, 312–316 (2020).

44. Carter, A.P. Crystal clear insights into how the dynein motor moves. Journal of cell science 126, 705–713 (2013).

45. Reimer, J.M., DeSantis, M.E., Reck-Peterson, S.L. & Leschziner, A.E. Structures of human dynein in complex with the lissencephaly 1 protein, LIS1. eLife 12, 84302 (2023).

46. Kon, T., Mogami, T., Ohkura, R., Nishiura, M. & Sutoh, K. ATP hydrolysis cycle-dependent tail motions in cytoplasmic dynein. Nature structural & molecular biology 12, 513–519 (2005).

47. Lee, W.L., Oberle, J.R. & Cooper, J.A. The role of the lissencephaly protein Pac1 during nuclear migration in budding yeast. J Cell Biol 160, 355–364 (2003).

48. Lenz, J.H., Schuchardt, I., Straube, A. & Steinberg, G. A dynein loading zone for retrograde endosome motility at microtubule plus-ends. The EMBO journal 25, 2275–2286 (2006).

49. Egan, M.J., Tan, K. & Reck-Peterson, S.L. Lis1 is an initiation factor for dynein-driven organelle transport. J Cell Biol 197, 971–982 (2012).

50. Jha, R., Roostalu, J., Cade, N.I., Trokter, M. & Surrey, T. Combinatorial regulation of the balance between dynein microtubule end accumulation and initiation of directed motility. The EMBO journal 36, 3387–3404 (2017).

51. Lammers, L.G. & Markus, S.M. The dynein cortical anchor Num1 activates dynein motility by relieving Pac1/LIS1-mediated inhibition. J Cell Biol 211, 309–322 (2015).

52. Ton, W.D. et al. Microtubule-binding-induced allostery triggers LIS1 dissociation from dynein prior to cargo transport. Nature structural & molecular biology, doi: 10.1038/s41594-41023-01010-x (2023).

53. Pecreaux, J. et al. Spindle oscillations during asymmetric cell division require a threshold number of active cortical force generators. Current biology 16, 2111–2122 (2006).

54. Yang, G. et al. Architectural dynamics of the meiotic spindle revealed by single-fluorophore imaging. Nature cell biology 9, 1233–1242 (2007).

55. Elshenawy, M.M. et al. Cargo adaptors regulate stepping and force generation of mammalian dynein-dynactin. Nat Chem Biol 15, 1093–1101 (2019).

